# Spatiotemporal Dynamics of Protein Recruitment During Cell Wound Repair

**DOI:** 10.64898/2026.08.14.744976

**Authors:** Mitsutoshi Nakamura, Justin Hui, Jeffrey M. Verboon, Susan M. Parkhurst

**Author notes:** Corresponding author: (SMP).

## Abstract

Injuries to individual cells happen frequently as a result of physiological and environmental stresses during their normal daily functions that can lead to a ruptured cell cortex (plasma membrane and underlying cortical cytoskeleton). The capacity of cells to rapidly repair general daily injuries, as well as ones resulting from trauma, infection, or diseases/cancer, is essential for their survival. While we know the general cell biological outline of the highly-conserved physiological events taking place during cell wound repair, our knowledge of the molecular mechanisms governing the repair process is still fairly limited, due in large part to the lack of information regarding the molecules, machineries, and pathways involved. Here, we performed a genetic screen of 1322 fluorescent-tagged proteins to identify cell wound repair components that are recruited upon laser wounding or whose expression is lost and/or altered upon laser wounding. We identified 129 proteins that are recruited to wounds during the cell repair process through high resolution spatio-temporal expression analyses of these gene fusions in conjunction with a fluorescent actin reporter. Strikingly, we find that many members of the Rab family GTPases are recruited to wounds where, in addition to their well-known roles in intracellular membrane trafficking, they are affecting actin cytoskeletal organization and dynamics during the repair process. These studies are allowing us to define the earliest acting proteins, as well as those required at specific steps in the repair process based on their recruitment patterns and the precise timing of their recruitment to wounds. Thus, our imaging-based screen is providing us with a global view of the repair processes, as well as a large number of genes/gene families that provide new entry points for examining specific steps in the cell wound repair process.

**Author Summary:** Cells in our bodies get injured every day from normal activity, environmental stress, infection, or disease. To survive, they must quickly repair these injuries and restore normal function. While some molecules have been identified as key players of cell wound repair, many of the molecules involved and their roles remain unknown. In this study, we identified new molecules that are involved in different steps of cell wound repair. Using laser-induced injury in the Drosophila model, we examined 1322 proteins and observed their spatial and temporal dynamics in a cell after injury. From the 1322 proteins examined, we identified 129 proteins recruited to distinct regions around the damage site during cell wound repair, suggesting roles in specific steps of the repair process. Interestingly, a subset of these proteins are Rab family GTPase members, highlighting new roles for these proteins in regulating actin dynamics. By identifying new candidate repair molecules, we provide a foundation for understanding how cells maintain their integrity and how repair processes may be influenced by factors such as wound size, infection, aging, and disease.

## Introduction

The plasticity of the cell cortex – plasma membrane and underlying cortical cytoskeleton – is vital to cellular functions, as well as when things go awry. Injuries to individual cells happen frequently as a result of daily wear-and-tear, accidents/trauma, violence, clinical procedures, and pathological conditions from infections to diseases and cancers (1–6). Cellular wounds can be particularly problematic when they occur in non-renewing or irreplaceable cell types, or occur together with fragile cell disease conditions such as those with muscular dystrophies, acute lung injuries, and diabetes. In addition, abnormal and/or excessive repair can induce or worsen physiological conditions such as fibrosis, inflammation, and tumorigenesis (1–3, 6). Damage to cells requires immediate sealing of the membrane breach, followed by wound closure then remodeling of the cell cortex to return it to its pre-wounded state. Wound repair shares many features with normal cellular and developmental events (4, 5), as well as with tumor progression (1–3, 7).

Cell wound repair is highly conserved and a number of complementary model systems have been developed to study this process, including the *Drosophila* syncytial embryo (8, 9), *Xenopus* oocytes (10, 11), *Dictyostelium* (12, 13), sea urchin eggs (14, 15), budding yeast (16, 17), and tissue culture cells (7, 18–22). While the details can vary among the different cell wound repair models, the physiological/cell biological sequence of events to close the wound following a breach to the cell cortex are similar (Fig 1A, 1 C) (3–5, 23). The first event following disruption of the cell cortex that sets the repair process in motion is a sudden influx of extracellular calcium. In response, the cell rapidly reseals the plasma membrane to prevent excessive loss of intracellular content by recruiting vesicles to the injury site that form a temporary membrane patch to plug the hole. Depending on the model system and wound size, the cell then either assembles an F-actin ring around the wound periphery that translocates inward to close the wound or excises the damaged cortex region using endocytosis/exocytosis. This is followed by a remodeling phase to remove repair structures (i.e., membrane plug and actomyosin ring remnants), and to re-establish the original cell cortex organization, connections, and functions.

**Figure 1.**
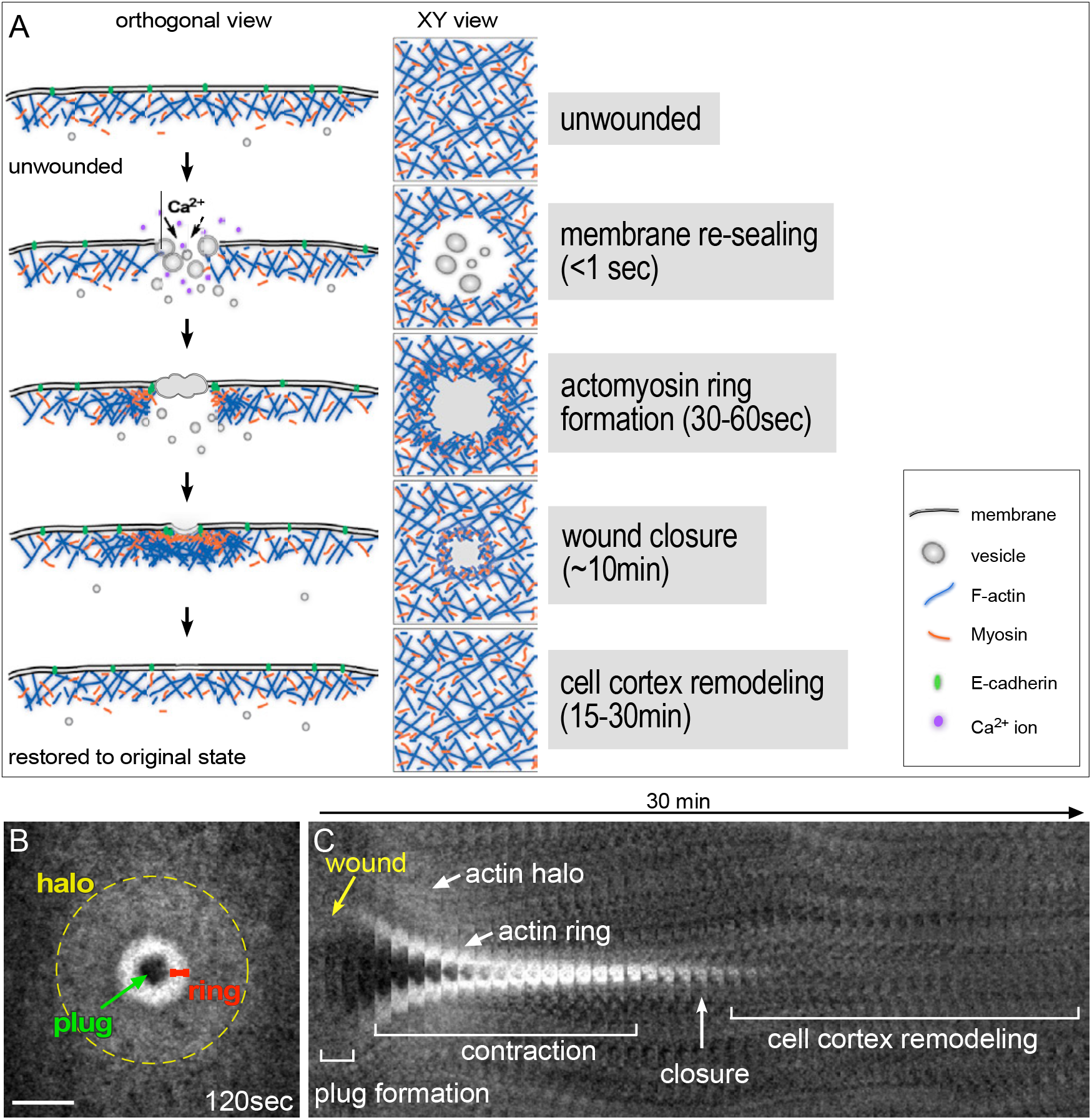
Cell Wound Repair. (A) Repair of cell cortex lesions begins with wound expansion, followed by the formation of a temporary membrane plug to re -seal the hole: intracellular membrane vesicles are recruited to the wound where they fuse with one another and the plasma membrane t o form this plug. After the membrane has been re -sealed, assembly and contraction of Rho family GTPase -dependent actomyosin arrays are used to close the cell cortex. After wound closure, re -establishing a normal cyto-architecture requires remodeling of the membrane and underlying cortical cytoskeleton to return it to its normal state. (B-C) Confocal xy projection (B) and kymograph (C) of a laser wound generated in a NC4 staged *Drosophila* embryo expressing a GFP-actin reporter. Actin accumulates in two regions: a highly enriched actin ring abutting the wound edge (red line), and an elevated actin halo encircling the actin ring (dashed circle). Membrane plug is marked with red arrow. Kymograph of wound repair showing the major features and events of cell wound repair. Scale bar: 20μm.

Our knowledge of the molecular mechanisms governing the cell wound repair process is still fairly limited, due largely to the lack of information regarding the molecules, machineries, and pathways involved. Here we use the *Drosophila* cell wound repair model that provides a powerful genetic model system with superb genetic resources and amenability for dynamic live imaging (8, 9, 24). Our earlier work established the early syncytial *Drosophila* embryo as a quantitatively robust model for studying cell wound repair. In this model, actin accumulates in two adjacent regions: 1) a highly-enriched ring at the wound edge, and 2) a less dense halo region encircling the actomyosin ring at the wound periphery (Fig 1B-C). Following these actin dynamics allows us to identify the different stages of the repair process, as well as to quantitatively assess the effect of knockdowns and/or mutants on this process. Our prior work has shown that protein recruitment to cell wounds is a strong indicator of a functional role in repair (cf. (8, 9, 24–30). Therefore, to identify cell wound repair components that are recruited upon laser wounding or whose expression is lost and/or altered upon laser wounding, we performed an imaging-based screen of 1322 Drosophila lines containing fluorescent-tagged proteins. We identified 129 proteins that are recruited to wounds during the cell repair process through high resolution spatio-temporal expression analyses of these gene fusions in conjunction with a fluorescent actin reporter to build a molecular pathway for cell wound repair. In addition to providing a global view of the repair processes, these studies are allowing us to define the earliest acting proteins, as well as those required at specific steps in the repair process based on their recruitment patterns and the precise timing of their recruitment to wounds. Thus, our imaging-based screen provided us with a global view of the repair processes, as well as a large number of genes/gene families that provide new entry points for examining specific steps in the cell wound repair process.

## Results

To identify proteins that are recruited to cell wounds or whose expression is lost and/or altered upon wounding, we performed a genetic screen of 1322 fluorescently-tagged proteins (Table S1, Table S2). We identified 129 proteins that are involved in the cell wound repair process (Table 1). The fluorescently-tagged proteins examined are from FlyTrap collections (31–36), as well as candidate proteins that have been generated by many labs and reside in the Bloomington, Kyoto, or Vienna Drosophila Stock Centers (Table S2). We determined the precise spatial recruitment patterns for these fluorescently-tagged proteins following wounding using time lapse spinning disk microscopy. Wounds were generated by laser ablation on the lateral side of nuclear cycle 4-6 *Drosophila* syncytial embryos expressing the fluorescently-tagged protein of interest, along with a fluorescent actin reporter (sChMCA, sGMCA, sK2MCA, or sStMCA; see Methods). Actin accumulates in a highly enriched actomyosin ring bordering the wound edge and an elevated actin halo encircling the actin ring, and serve as reference point for the localization of the screened proteins (Fig 1B-C). The patterns of protein recruitment could be divided into 7 broad categories: plug region, a ring inside the actin ring, a ring overlapping the actin ring, actin ring plus actin halo, actin halo, multiple recruitment patterns, and all other patterns. For each protein recruited, we show kymographs of the fluorescently-tagged protein with and without the actin reporter, as well as the split and merged channels of an appropriate time point for that recruitment pattern with a line plot demonstrating the spatial localization of the protein in relation to the actin reporter. Some of the fluorescently-tagged proteins examined were hard to image due to their low expression, transient expression, propensity to bleach quickly during the time lapse, and/or the interference of non-wound fluorescent expression (i.e., high endoplasmic reticulum expression). For these cases, we show a single time point and line plot to document their recruitment to cell wounds.

**Table 1.**
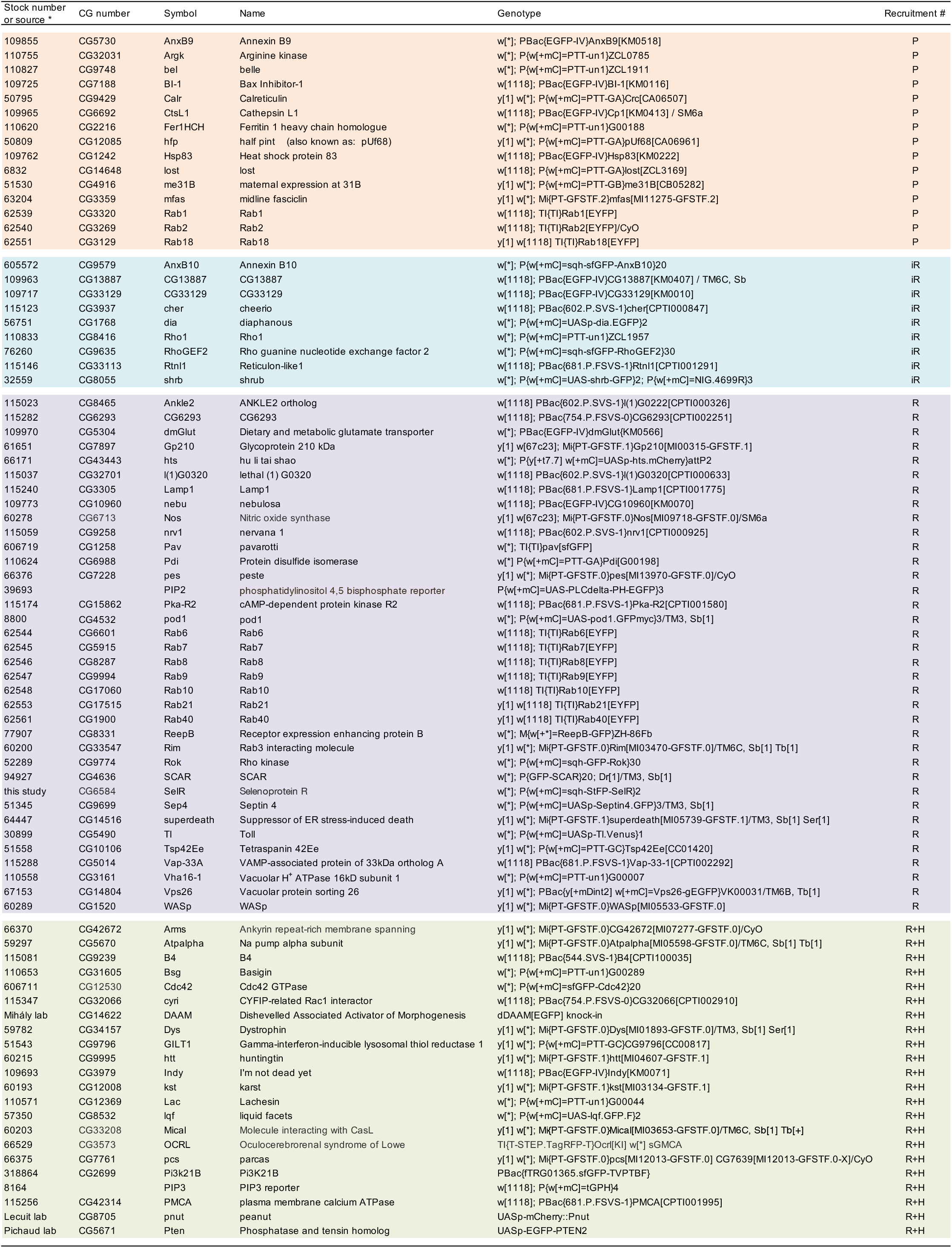
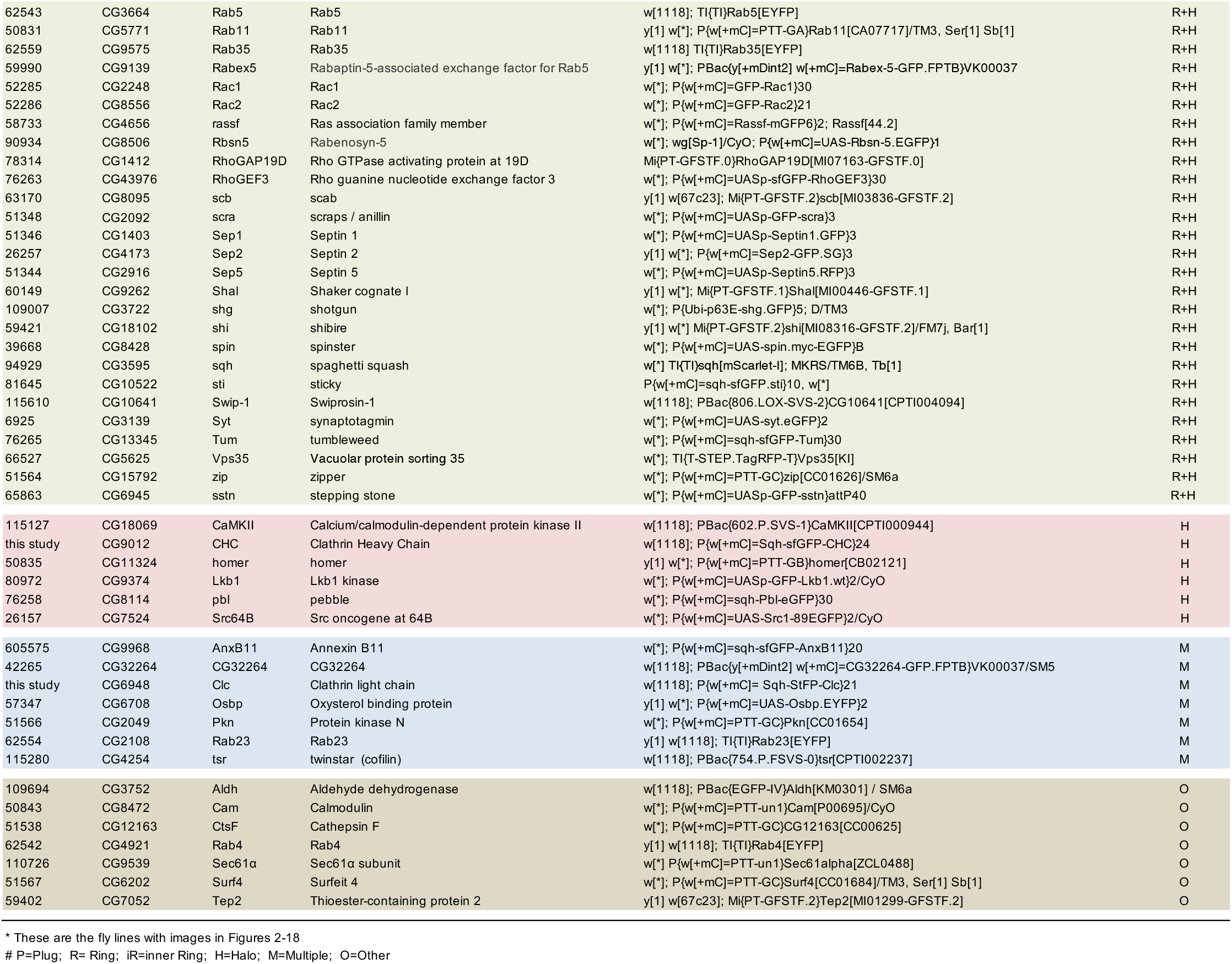
Fluorescently-tagged proteins recruited to cell wounds.

### Proteins recruited to the plug region of cell wounds

The region inside of the wound (plug region) houses a temporary membrane plug that is formed rapidly after wounding to stop cytoplasmic loss and to block the entry of extracellular material. We identified 12 proteins that are recruited to the central-most part of the plug region inside the wound: AnxB9, Argk, Bel, Bl-1, Calr, CtsL-1, Fer1HCH, Hfp, Hsp83, Lost, Me31B, and Mfas (Fig 2; Table 1). In most cases, this recruitment to the center of the wound was accompanied by a concentric ring of protein recruitment exclusion. The exceptions to this are Bl-1, Fer1HCH, and Mfas where their protein recruitment fills the area inside the actin ring (Figs 2I-J, 2L, 2U-V, 2X). There is not an obvious common theme that ties these different proteins together, i.e., they are not all membrane associated proteins as might be expected as they are recruited to the membrane plug region inside the wound. Our previous studies showed that one of these, AnxB9, a calcium responsive protein, is recruited very rapidly (<3 sec) where it stabilizes actin to regulate the subsequent recruitment of RhoGEF2 to form a robust actomyosin ring (9).

**Figure 2.**
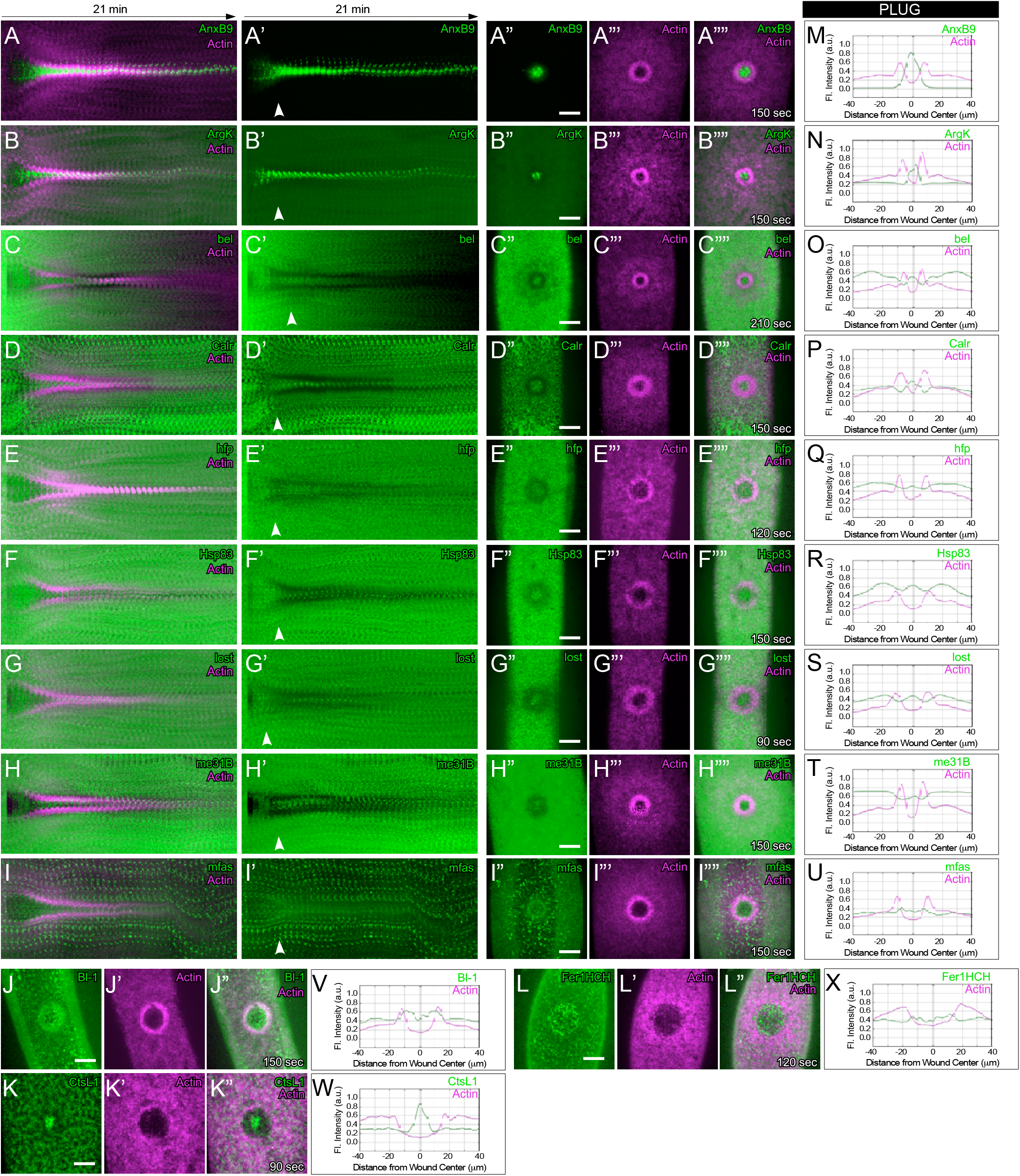
Proteins recruited to the plug region of cell wounds. (A-L”) Confocal max projection images of embryos expressing a fluorescent actin reporter (sGMCA or sStMCA) and the following fluorescently-tagged proteins: AnxB9 (A-A””), Argk (B-B””), bel (C-C””), Calr (D-D””), hfp (E-E””), Hsp83 (F-F””), lost (G-G””), me31B (H-H””), mfas (I-I””), Bl-1 (J-J”), CstL1 (K-K”), or Fer1HCH (L-L”), at the time points indicated. (A -I’) Kymographs across the wound area in A -I””, respectively. White arrows indicate the time point of the XY views shown in A” -I””. (A”-I””, J-L”) XY views of the time point indicated across the wound area in A -L”, respectively. (M-X) Fluorescence intensity (arbitrary units) profiles across the wound area over time for the images shown in (A”” -I””, J”-L”). Scale bars: 20μm.

### Proteins recruited to the wound edge just inside the actin ring region

The wound edge is a key assembly site for the molecular machineries required to integrate polymerized actin into the actin ring, as well as the fusion site between the temporary membrane plug and the intact plasma membrane. We identified many proteins that are recruited to the wound periphery. Interestingly, for 9 of these, the ring that is formed lies <u>internal</u> to the actin ring (“inner ring”): AnxB10, CG13387, CG33129, Cher, Dia, Rho1, RhoGEF2, Rtnl-1, and Shrb (Fig 3; Table 1). AnxB10, another calcium responsive protein, is also recruited very rapidly (<3 sec) where it regulates actin dynamics that affect the subsequent recruitment of RhoGEF3 to the actin ring region (29). Previous work has shown that Rho family GTPases form concentric rings at the wound periphery with Rho1 being inside of the actin ring (9, 25). RhoGEF2, Rho1, and Dia are also known to be part of the molecular pathway needed to form the contractile actomyosin ring that encircles the wound periphery and translocates inward to close the wound (9, 25). Knockdown of these genes results in wound over-expansion, lower actin accumulation at the actin ring, decreased actin ring width, and slow wound repair. While these phenotypes are also observed with AnxB10 knockdowns, additional phenotypes such as actin accumulation within the wound also occur (29).

**Figure 3.**
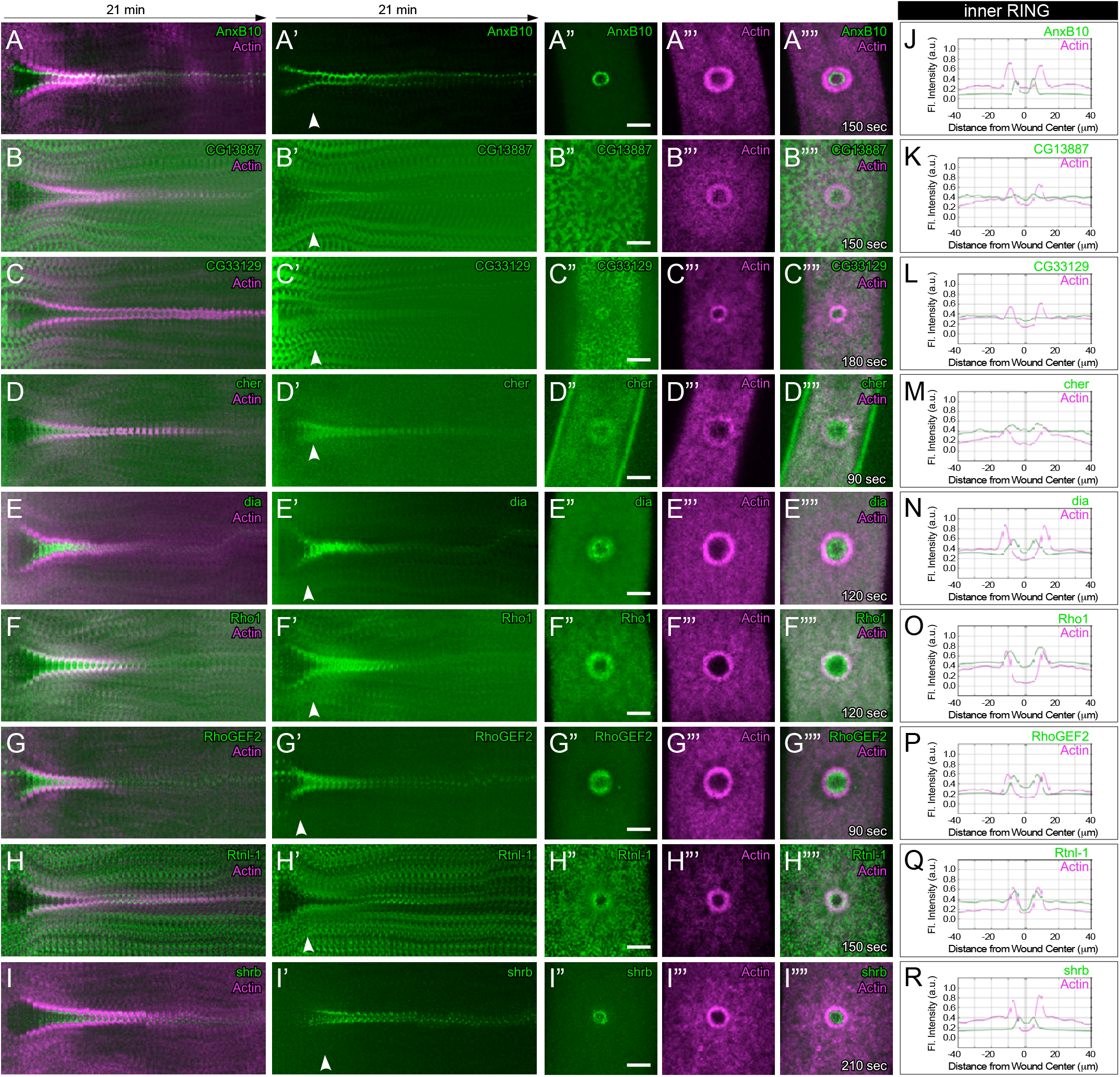
Proteins recruited to the wound edge just inside the actin ring region. (A-I””) Confocal max projection images of embryos expressing a fluorescent actin reporter (sGMCA or sStMCA) and the following fluorescently-tagged proteins: AnxB10 (A-A””), CG13887 (B-B””), CG33129 (C-C””), cher (D-D””), dia (E-E””), Rho1 (F-F””), RhoGEF2 (G-G””), Rtnl-1 (H-H””), or shrb (I-I””), at the time points indicated. (A - I’) Kymographs across the wound area in A -I””, respectively. White arrows indicate the time point of the XY views shown in A”-I””. (A”-I””) XY views of the time point indicated across the wound area in A -I””, respectively. (J-R) Fluorescence intensity (arbitrary units) profiles across the wound area over time for the images shown in (A””-I””). Scale bars: 20μm.

### Recruited proteins overlapping the actin ring region of cell wounds

The formation of a robust actin ring at the wound periphery is required to generate the physical force necessary to pull the cell cortex inward for wound closure. Many proteins that are recruited to the wound periphery overlap with the actin ring. We identified 29 proteins with this recruitment pattern (Figs 4-6). The ectopic expression of some of these fluorescently-tagged proteins disrupts the actin reporter (cf. hts, ReepB, Rim) suggesting a role for these proteins in actin ring formation/organization, or disrupts wound closure dynamics (cf. dmGlut, Nos, Pod1) suggesting a role in actin dynamics. In addition, knockdown of four of these genes (Pav, SCAR, Sep4, and WASp) have been shown to affect actin dynamics, including actin bending and actomyosin ring assembly, resulting in impaired cell wound repair phenotypes (24, 27, 28, 30).

**Figure 4.**
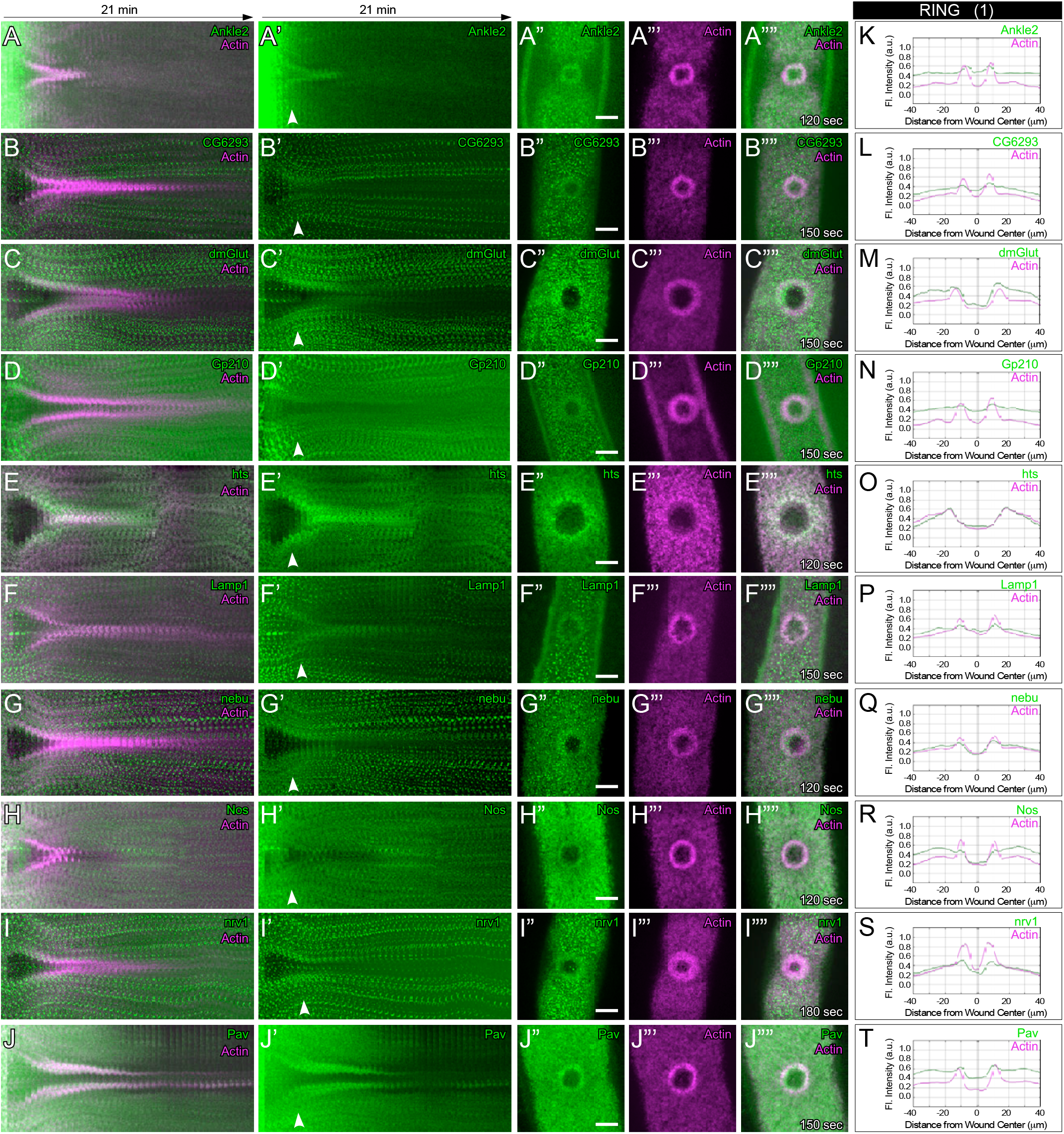
Recruited proteins overlapping the actin ring region of cell wounds (1). (A-J””) Confocal max projection images of embryos expressing a fluorescent actin reporter (sGMCA or sStMCA) and the following fluorescently-tagged proteins: Ankle2 (A-A””), CG6293 (B-B””), dmGlut (C-C””), Gp210 (D-D””), hts (E-E””), Lamp1 (F-F””), nebu (G-G””), Nos (H-H””), nrv1 (I-I””), or Pav (J-J””), at the time points indicated. (A-J’) Kymographs across the wound area in A -J””, respectively. White arrows indicate the time point of the XY views shown in A” -J””. (A”-J””) XY views of the time point indicated across the wound area in A-J””, respectively. (K-T) Fluorescence intensity (arbitrary units) profiles across the wound area over time for the images shown in (A”” -J””). Scale bars: 20μm.

**Figure 5.**
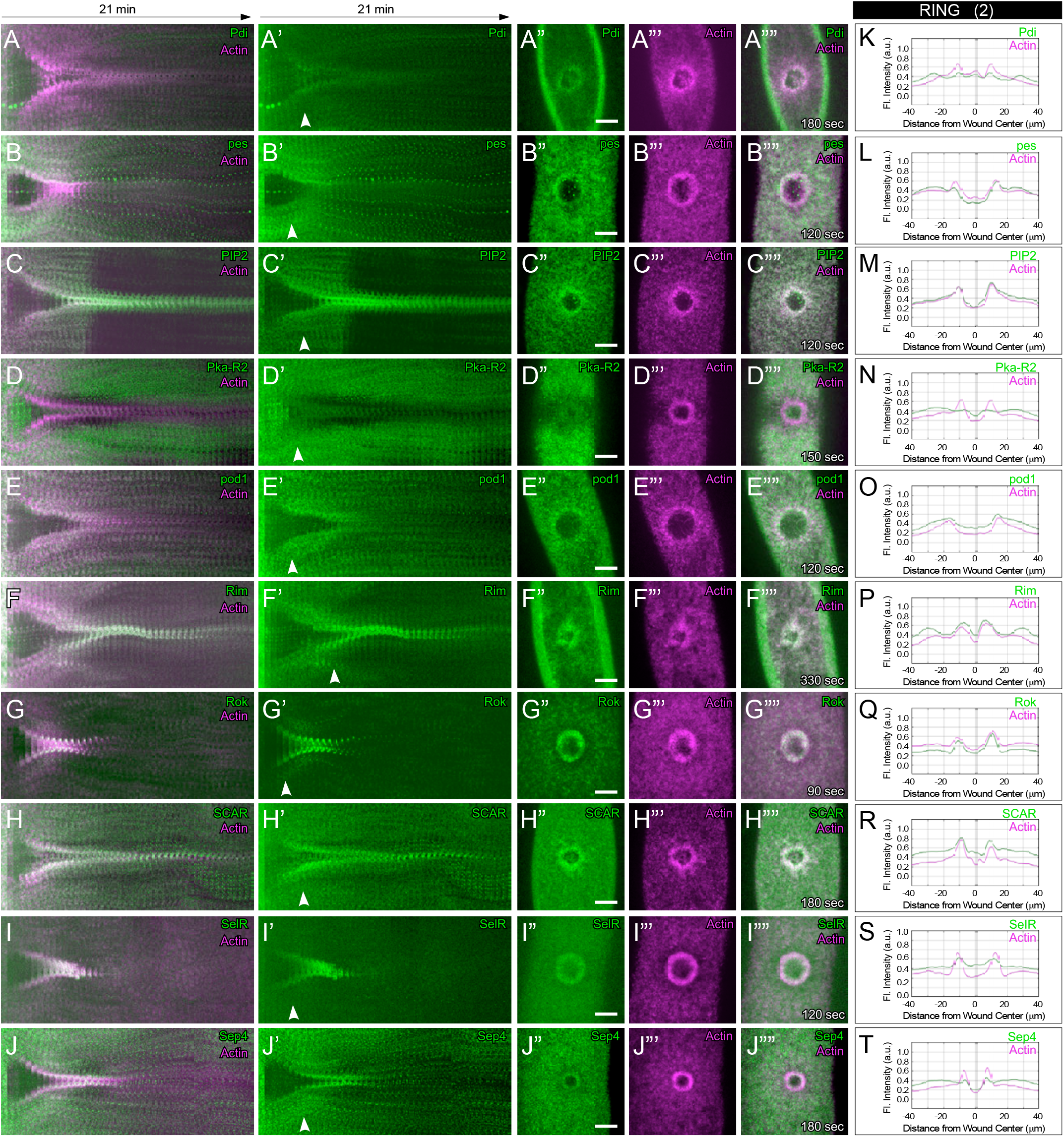
Recruited proteins overlapping the actin ring region of cell wounds (2). (A-J””) Confocal max projection images of embryos expressing a fluorescent actin reporter (sGMCA or sStMCA) and the following fluorescently-tagged proteins: Pdi (A-A””), pes (B-B””), PIP2 (C-C””), Pka-R2 (D-D””), pod1 (E-E””), Rim (F-F””), Rok (G-F””), SCAR (H-G””), SelR (I-I””), or Sep4 (J-J””), at the time points indicated. (A - J’) Kymographs across the wound area in A -J””, respectively. White arrows indicate the time point of the XY views shown in A”-J””. (A”-J””) XY views of the time point indicated across the wound area in A -J””, respectively. (K-T) Fluorescence intensity (arbitrary units) profiles across the wound area over time for the images shown in (A””-J””). Scale bars: 20μm.

**Figure 6.**
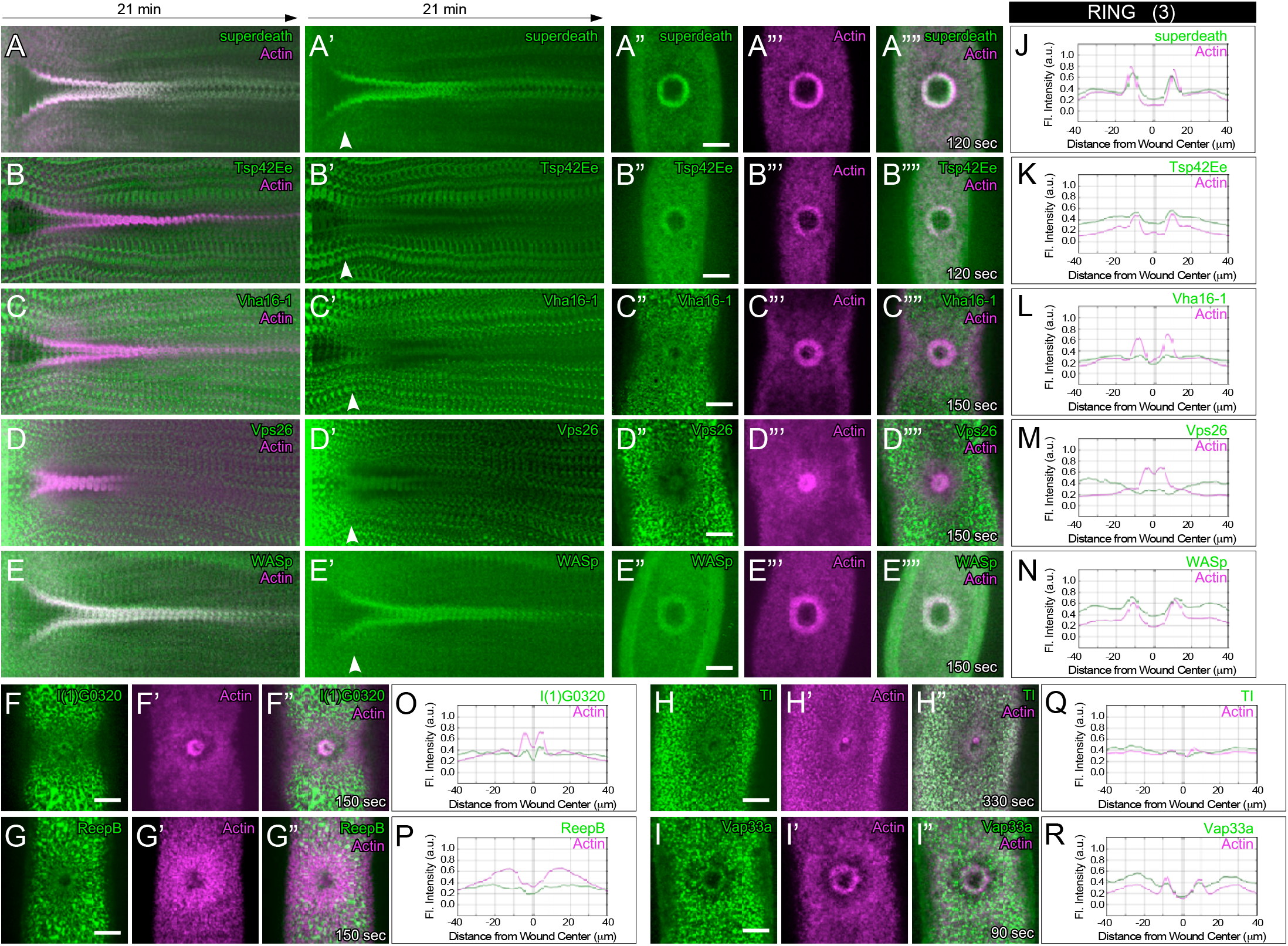
Recruited proteins overlapping the actin ring region of cell wounds (3). (A-I”) Confocal max projection images of embryos expressing a fluorescent actin reporter (sGMCA or sStMCA) and the following fluorescently-tagged proteins: superdeath (A-A””), Tsp42Ee (B-B””), Vha16-1 (C-C””), Vps26 (D-D””), WASp (E-E””), l(1)G0320 (F-F””), ReepB (G-G””), Tl (H-H””), or Vap33a (I-I””), at the time points indicated. (A-E’) Kymographs across the wound area in A-E””, respectively. White arrows indicate the time point of the XY views shown in A”-E””. (A”-E ””) XY views of the time point indicated across the wound area in A-E”, respectively. (J-R) Fluorescence intensity (arbitrary units) profiles across the wound area over time for the images shown in (A””-E””, F”-I”), respectively. Scale bars: 20μm.

### Recruited proteins overlapping the actin ring and halo regions of cell wounds

In addition to the immediate wound periphery where the dense actin ring forms, a region of less dense actin recruitment surrounds this actin ring (actin halo; Fig 1B). Protein recruitment to wounds that overlaps both the actin ring and actin halo is the largest category of proteins with 46 of these identified (Figs 7-11; Table 1). This category is similar to those that overlap with the actin ring only: they encode a wide variety of activities/functions, and the ectopic expression of some of these affects the actin reporter or wound closure dynamics (cf. scb, sstn). Knockdown of several of these genes (Cdc42, DAAM, PIP3, pnut, Rac1, RhoGEF3, scra, Sep1, Sep2, Sep5, shg, sqh, Tum, and Zip) has been shown to affect actin dynamics resulting in a variety of impaired cell wound repair phenotypes (8, 9, 24, 25, 27, 28, 30).

**Figure 7.**
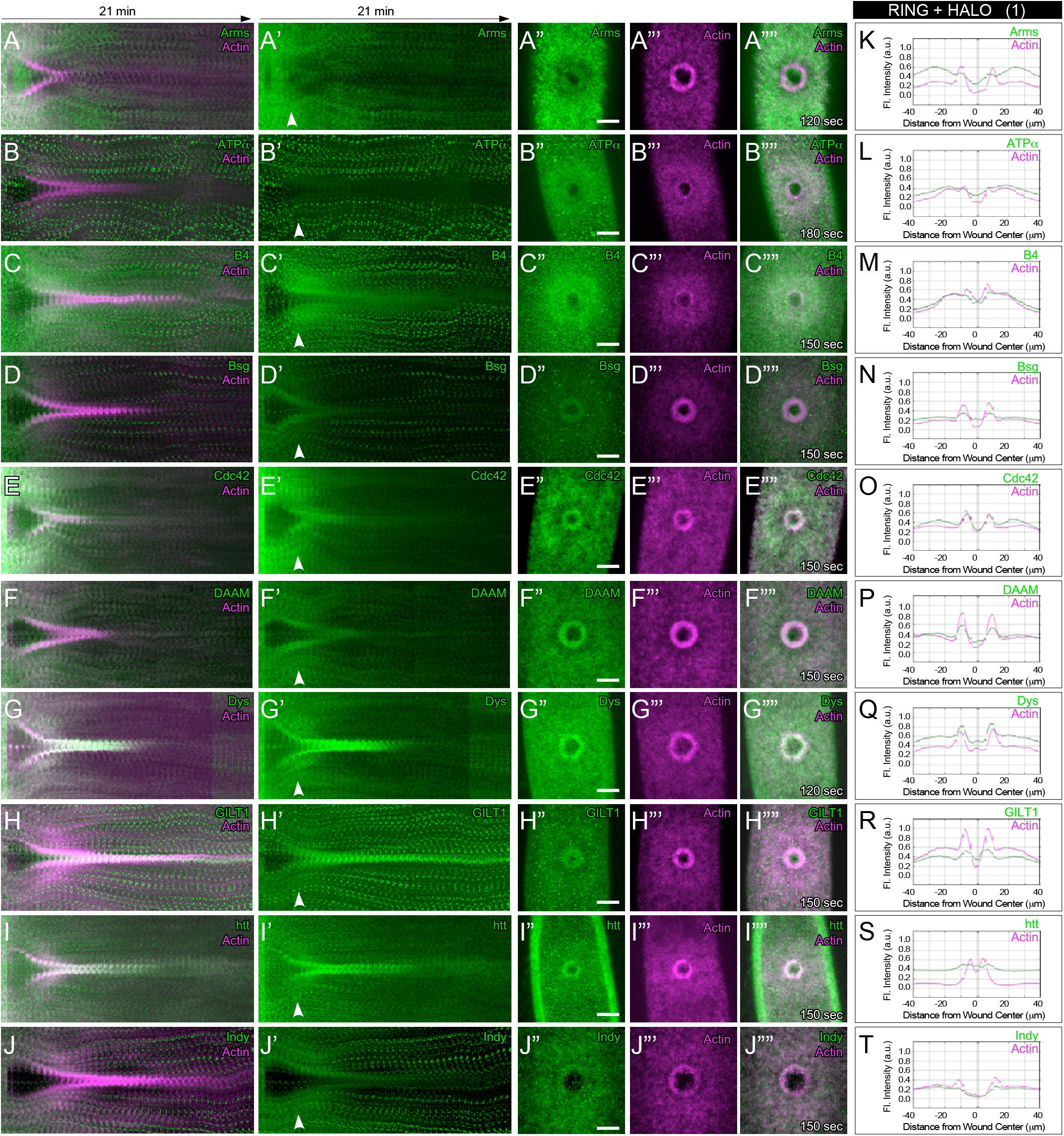
Recruited proteins overlapping the actin ring and halo regions of cell wounds (1). (A-J””) Confocal max projection images of embryos expressing a fluorescent actin reporter (sGMCA or sStMCA) and the following fluorescently-tagged proteins: Arms (A-A””), ATPα (B-B””), B4 (C-C””), Bsg (D-D””), Cdc42 (E-E””), DAAM (F-F””), Dys (G-G””), GILT1 (H-H””), htt (I-I””), or Indy (J-J””), at the time points indicated. (A-J’) Kymographs across the wound area in A -J””, respectively. White arrows indicate the time point of the XY views shown in A” -J””. (A”-J””) XY views of the time point indicated across the wound area in A-J””, respectively. (K-T) Fluorescence intensity (arbitrary units) profiles across the wound area over time for the images shown in (A”” -J””). Scale bars: 20μm.

**Figure 8.**
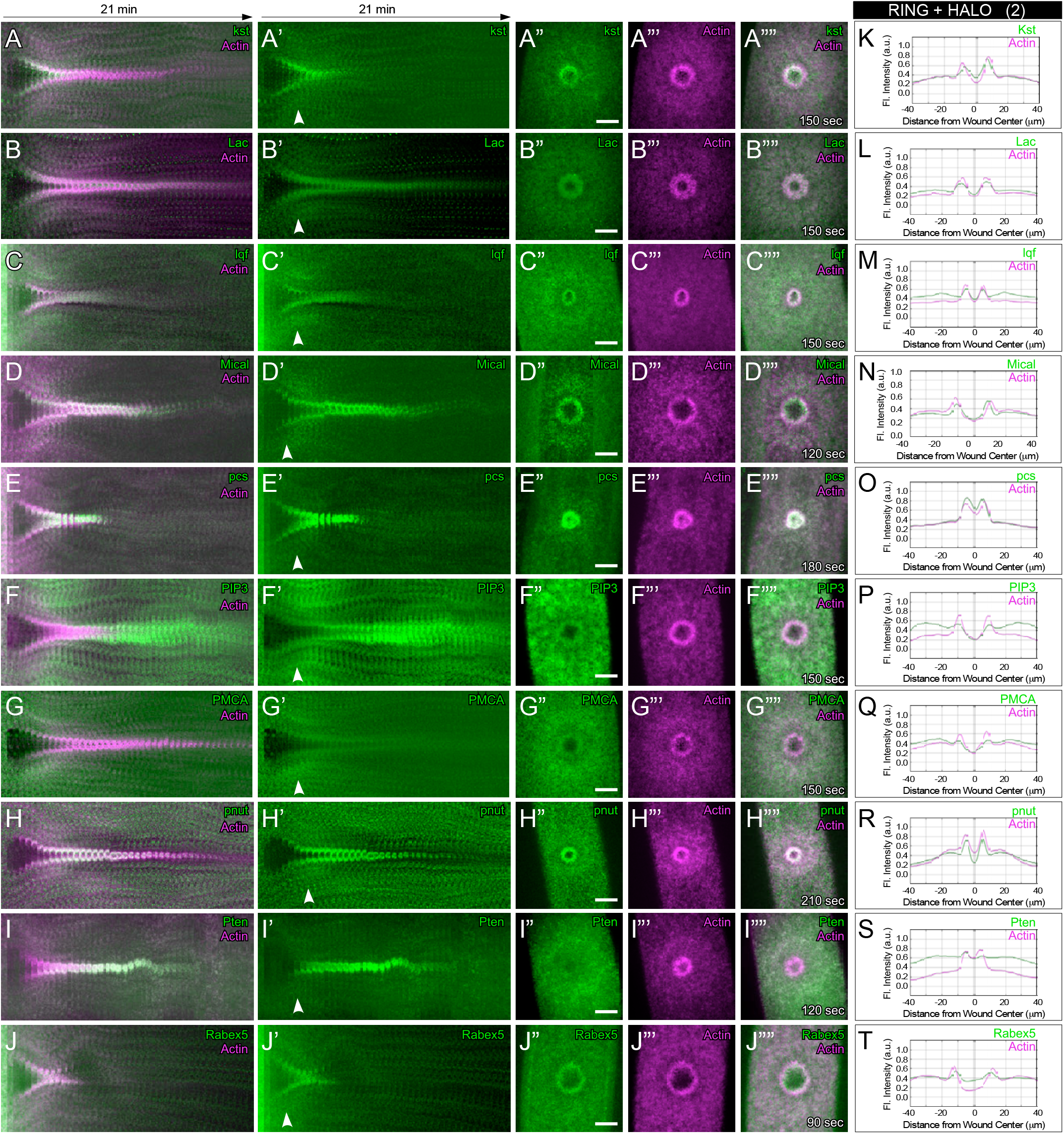
Recruited proteins overlapping the actin ring and halo regions of cell wounds (2). (A-J””) Confocal max projection images of embryos expressing a fluorescent actin reporter (sGMCA or sStMCA) and the following fluorescently -tagged proteins: kst (A-A””), Lac (B-B””), lqf (C-C””), Mical (D-D””), pcs (E-E””), PIP3 (F-F””), PMCA (G-G””), pnut (H-H””), Pten (I-I””), or Rabex5 (J-J””), at the time points indicated. (A-J’) Kymographs across the wound area in A -J””, respectively. White arrows indicate the time point of the XY views shown in A”-J””. (A”-J””) XY views of the time point indicated across the wound area in A -J””, respectively. (K-T) Fluorescence intensity (arbitrary units) profiles across the wound area over time for the images shown in (A””-J””). Scale bars: 20μm.

**Figure 9.**
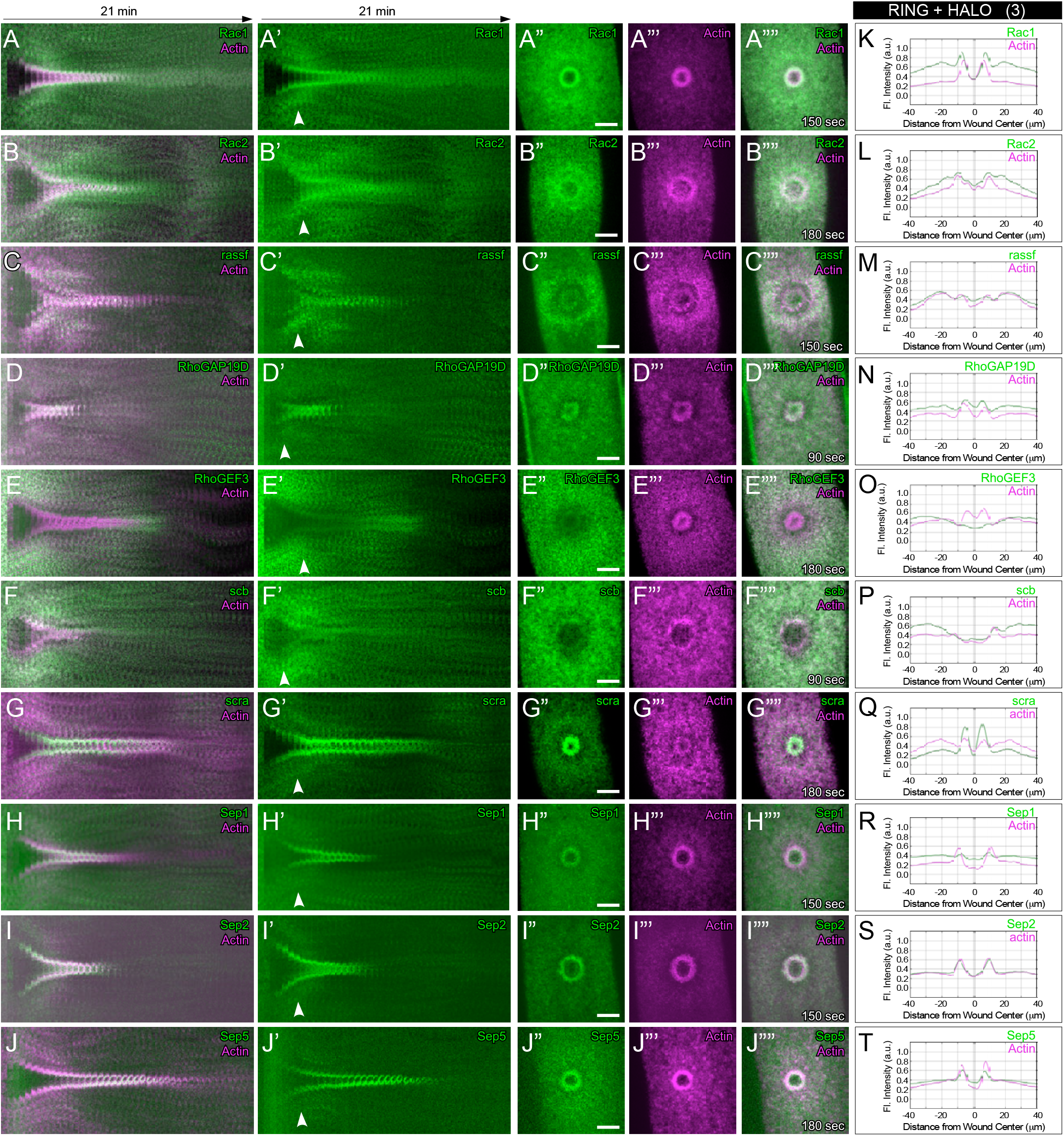
Recruited proteins overlapping the actin ring and halo regions of cell wounds (3). (A-J””) Confocal max projection images of embryos expressing a fluorescent actin reporter (sGMCA or sStMCA) and the following fluorescently-tagged proteins: Rac1 (A-A””), or Rac2 (B-B””), rassf (C-C””), RhoGAP19D (D-D””), RhoGEF3 (E-E””), scb (F-F””), scra (G-G””), Sep1 (H-H””), Sep2 (I-I””), or Sep5 (J-J””), at the time points indicated. (A-J’) Kymographs across the wound area in A -J””, respectively. White arrows indicate the time point of the XY views shown in A” -J””. (A”-J””) XY views of the time point indicated across the wound area in A-J””, respectively. (L-T) Fluorescence intensity (arbitrary units) profiles across the wound area over time for the images shown in (A”” -J””). Scale bars: 20μm.

**Figure 10.**
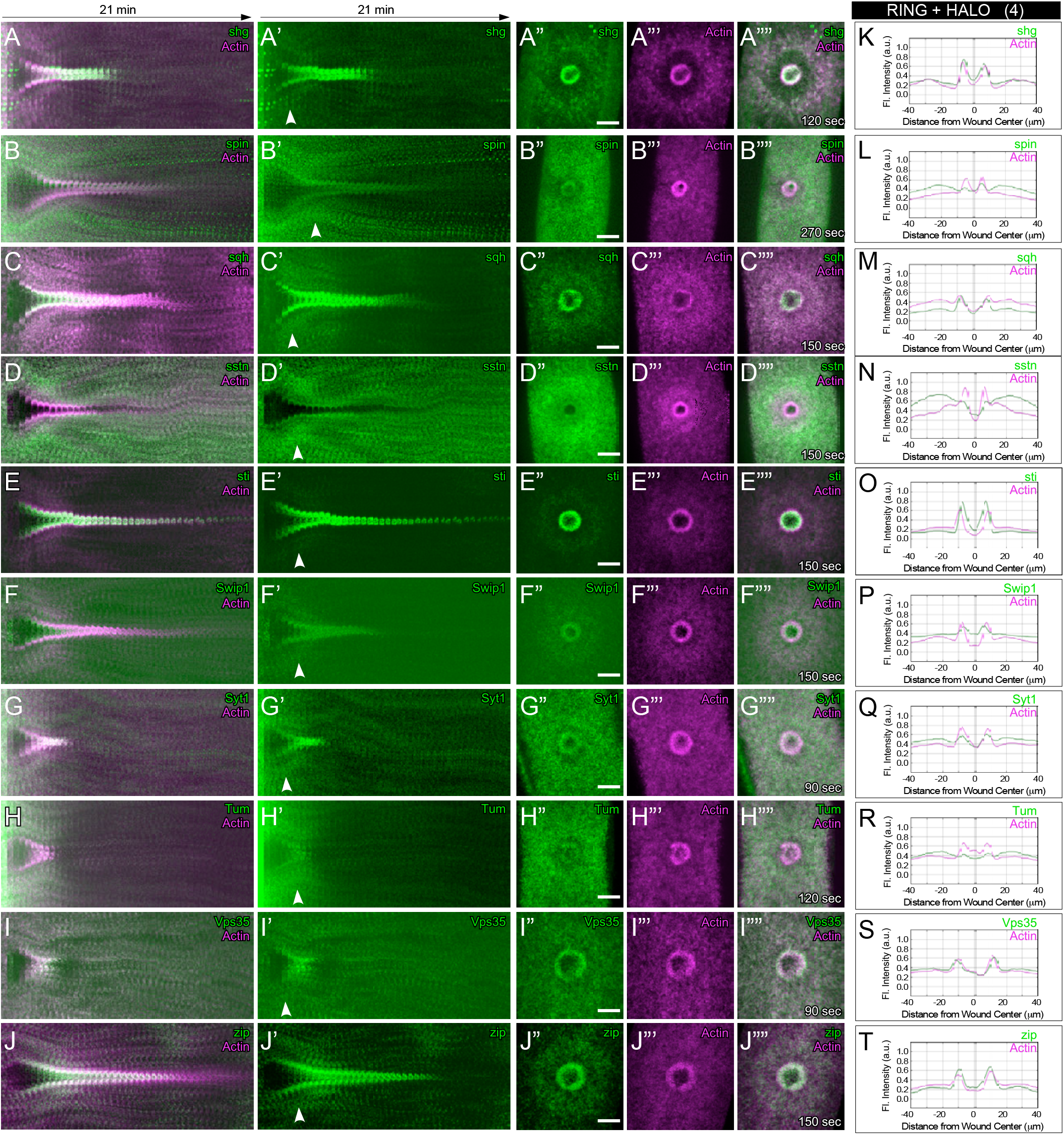
Recruited proteins overlapping the actin ring and halo regions of cell wounds (4). (A-J”) Confocal max projection images of embryos expressing a fluorescent actin reporter (sGMCA or sStMCA) and the following fluorescently -tagged proteins: shg (A-A””), spin (B-B””), sqh (C-C””), sstn (D-D””), sti (E-E””), Swip1 (F-F”), Syt1 (G-G””), Tum (H-H””), Vps35 (I-I””), or zip (J-J””), at the time points indicated. (A-J’) Kymographs across the wound area in A -J””, respectively. White arrows indicate the time point of the XY views shown in A”-J””. (A”-J””) XY views of the time point indicated across the wound area in A -J”, respectively. (K-T) Fluorescence intensity (arbitrary units) profiles across the wound area over time for the images shown in (A””-J””). Scale bars: 20μm.

**Figure 11.**
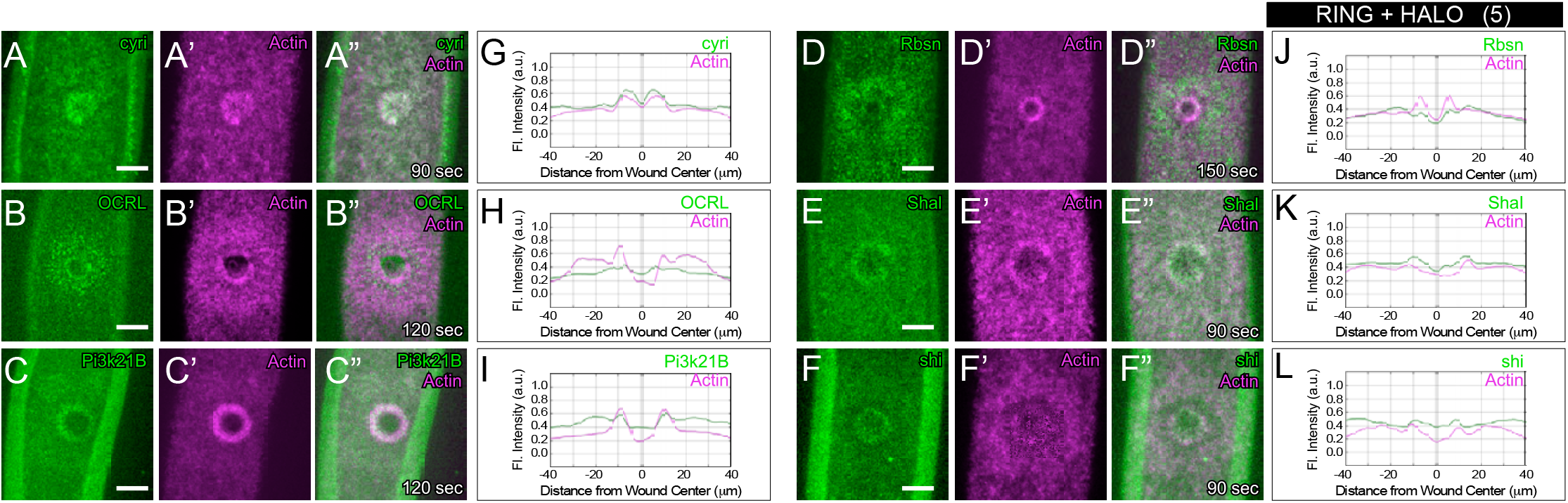
Recruited proteins overlapping the actin ring and halo regions of cell wounds (5). (A-F”) Confocal max projection XY images of embryos expressing a fluorescent actin reporter (sGMCA or sStMCA) and the following fluorescently-tagged proteins: cyri (A-A””), OCRL (B-B””), PI3k21B (C-C”), Rbsn (D-D”), Shal (E-E”), or shi (F-F”), at the time points indicated. (G-L) Fluorescence intensity (arbitrary units) profiles across the wound area over time for the images shown in (A” -F”). Scale bars: 20μm.

### Recruited proteins overlapping the actin halo region of cell wounds

While less is known about the role of the actin halo region in the repair process, it has been proposed to regulate the movement of cortical actin (and other repair components) towards the wound edge. We identified 7 proteins that are recruited to the actin halo region outside the wound: CaMKII, CHC, Homer, Lkb1, Pbl, and Src64B (Fig 12; Table 1). Of these, only the role of Pbl has been examined in cell wound repair to date. Knocking down Pbl results in severe cell wound repair phenotypes, including wound overexpansion, delayed actin accumulation around the wound edge, accumulation of actin inside the wound, and delayed repair (Fig 12E-E””, 12K) (9). This region could contribute to cell cortex stability/remodeling and further interrogation of the newly identified proteins localizing to this region may provide a definitive function.

**Figure 12.**
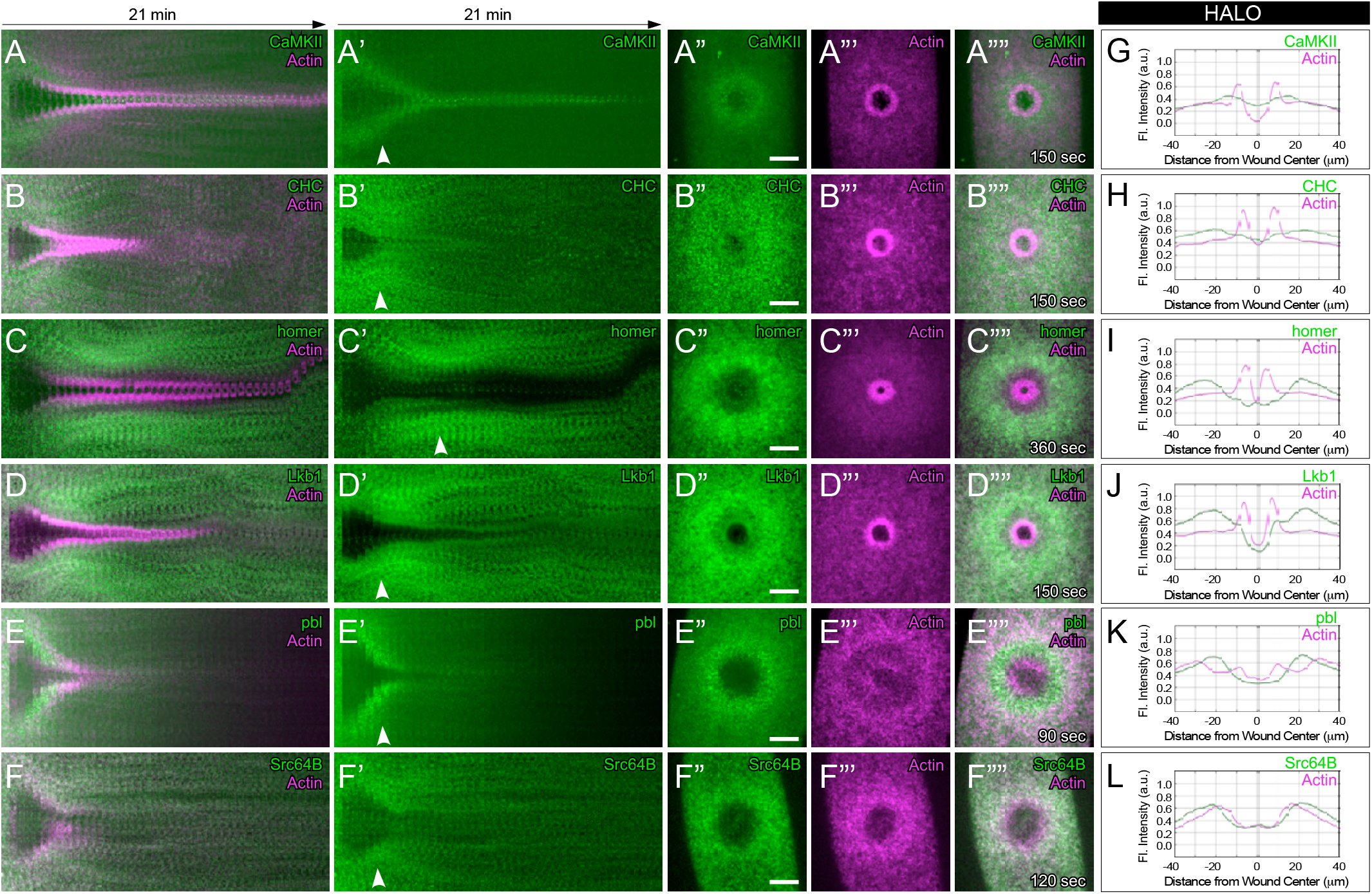
Recruited proteins overlapping the actin halo region of cell wounds. (A-F””) Confocal max projection images of embryos expressing a fluorescent actin reporter (sGMCA or sStMCA) and the following fluorescently-tagged proteins: CaMKII (A -A””), CHC (B-B””), homer (C-C””), Lkb1 (D-D””), pbl (E-E””), or Src64B (F-F””), at the time points indicated. (A -F’) Kymographs across the wound area in A -F””, respectively. White arrows indicate the time point of the XY views shown in A” -F””. (A”-F””) XY views of the time point indicated across the wound area in A -F””, respectively. (G-L) Fluorescence intensity (arbitrary units) profiles across the wound area over time for the images shown in (A”-F””). Scale bars: 20μm.

### Proteins exhibiting multiple recruitment patterns to cell wounds

Five of the proteins we examined exhibited dynamic recruitments to cell wounds with two distinct spatial recruitment patterns separated temporally: AnxB11, CG32264, Clc, Osbp, Pkn and tsr (Fig 13, Table 1). AnxB11 is a calcium responsive protein that is recruited to cell wounds rapidly (<3sec) (29). This robust recruitment overlaps with that of the actin ring (Fig 13Aiii-v, 13G). This AnxB11 recruitment then quickly collapses into the center of the wound where it fills the membrane plug area inside of the actin ring (Fig 13Avi-viii, 13G’). CG32264 is predicted to enable actin binding and affect actin cytoskeleton organization (37, 38). Its human orthologs, phosphatase and actin regulators (PHACTR1, −2, −4), are implicated in a variety of disease conditions, including multiple sclerosis, arterial diseases, and neurodegenerative diseases (Parkinsons) (37, 39–43). CG32264 is first recruited to the halo region (Fig 13Biii-v, 13H). Once the actin ring has closed, CG32264 accumulates in the center of the wound overlapping the actin ring (Fig 13Bvi-viii, 13H’). Clc (Clathrin light chain) is a major coating component of coated vesicles that is initially recruited to the actin halo region around the wound (Fig 13Ciii-v, 13I). As the wound approaches closure, this recruitment coalesces to overlap with the actin ring (Fig 13Cvi-viii, 13I’). The lipid binding protein Osbp (oxysterol binding protein) is initially recruited to cell wounds as a ring that lies just inside the actin ring (inner ring) (Fig 13Diii-v, 13J). As the actomyosin ring contracts to close the wound, the ring of Osbp moves such that it comes to lie outside of the actin ring (Fig 13Dvi-viii, 13J’). This Osbp recruitment dissipates as the actin ring is disassembled (Fig 13Di-ii). Protein kinase N (Pkn), a protein kinase that is activated by Rho family GTPases, shows a recruitment pattern similar to AnxB11: it is initially recruited to the region overlapping the actin ring (Fig 13 Eiii-v, 13K) and then collapses to the center of the wound in the membrane plug region (Fig 13Evi-viii, 13IK’). Twinstar (tsr), the Drosophila cofilin ortholog, encodes F-actin depolymerizing activity that enhances filament flexibility and facilitates filament severing (44–48). Tsr is initially recruited to the membrane plug region and is excluded from the actin ring region (Fig 13Fiii-v, 13L), where it could be working to clear random actin from the center of the wound. Several minutes later, it forms a ring encircling the nearly closed actin ring where the actin halo initially resides (Fig 13Fvi-viii, 13L’). This localization is consistent with a role in helping to disassemble the actin ring.

**Figure 13.**
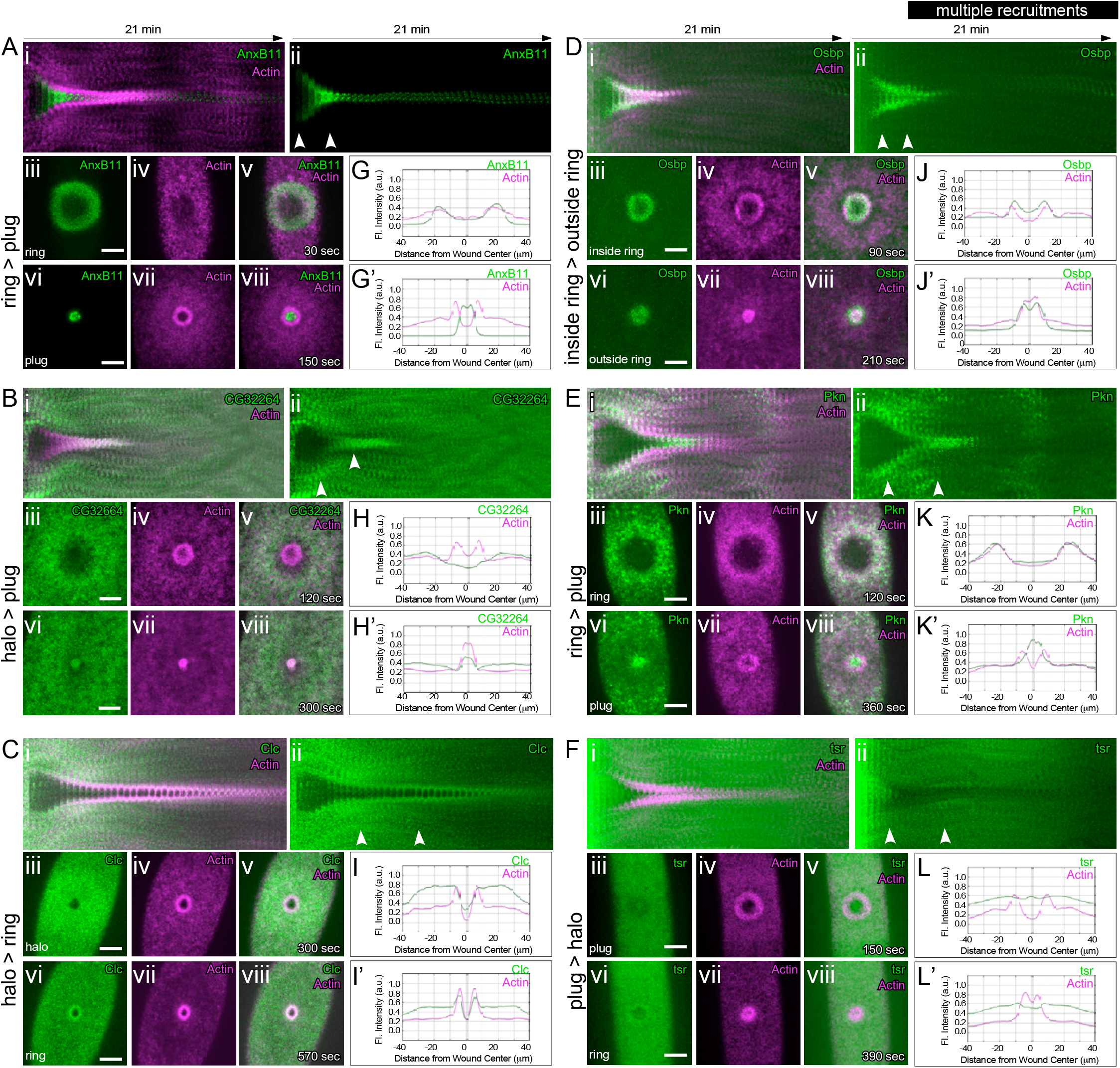
Proteins exhibiting multiple recruitment patterns to cell wounds. (A-E) Confocal max projection images of embryos expressing a fluorescent actin reporter (sGMCA or sStMCA) and the following fluorescently-tagged proteins: AnxB11 (A -Aviii), CG32264 (B-Bviii), Clc (C-Cviii), Osbp (D-Dviii), Pkn (E-Eviii), or tsr (F-Fviii), at the time points indicated. (Ai -Fii) Kymographs across the wound area in A - E, respectively. White arrows indicate the time point of the XY views shown in Aiii -Fviii””. (Aiii-Fviii) XY views of the time point indicated across the wound area in Aiii -Fviii, respectively. (G-L’) Fluorescence intensity (arbitrary units) profiles across the wound area over time for the images shown in (Aiii -Fviii). Scale bars: 20μm.

### Proteins exhibiting other recruitment patterns to cell wounds

A few of the proteins that we examined exhibited recruitment patterns that did not fall easily into the previous broad categories: Aldh, Cam, Sec61α, Surf4, CtsF, and Tep2 (Fig 14; Table 1). Aldh (Aldehyde dehydrogenase) exhibits a burst of robust recruitment to the wound area covering the actin halo region and a second concentric ring of recruitment outside of the actin halo region (“outer halo”) (Fig 14A-A’”, 14G). Aldh is known to detoxify aldehydes generated by lipid peroxidation (49, 50), it may be helping to modify and remodel the membrane during repair. Calmodulin (Cam), a Calcium-binding messenger protein, also exhibits a burst of robust recruitment to the entire wound area covering the plug, actin ring, actin halo and outer halo regions (Fig 14B-B’”, 14H). This Cam recruitment lasts ∼7 minutes (Fig 14B-B’), which is the time it takes for the initial influx of Calcium into wounds to dissipate (29). Sec61α encodes a subunit of translocon channels that function in the endoplasmic reticulum. Sec61α is rapidly recruited to the plug and actin ring regions (Fig 14C-C’”, 14I), where it may aide in forming the membrane plug and/or in the interaction on the plasma membrane with the underlying cortical cytoskeleton. Surf4 encodes an endoplasmic reticulum cargo receptor that mediates the export of lipoproteins. Surf4 does not appear to be recruited to the wound edge, but rather undergoes a redistribution such that it moves from the endoplasmic reticulum to a ring at the wound periphery (Fig 14D-D’”, 14J). CtsF and Tep2 are recruited to the wound periphery (Fig 14E-F’”, 14K-L), but both are unique in that this recruitment is extracellular (Fig 14E’”, 14F’”). CtsF encodes the Drosophila ortholog of Cathepsin F, a secreted lysosomal cysteine protease known to play a role in many processes including protein degradation (51–53). Tep2 encodes an endopeptidase inhibitor that is known to reside in extracellular spaces and function in defense responses (54–58). Its recruitment to the extracellular space outside cell wounds may allow it to protect the cell from bacterial or other extracellular insults.

**Figure 14.**
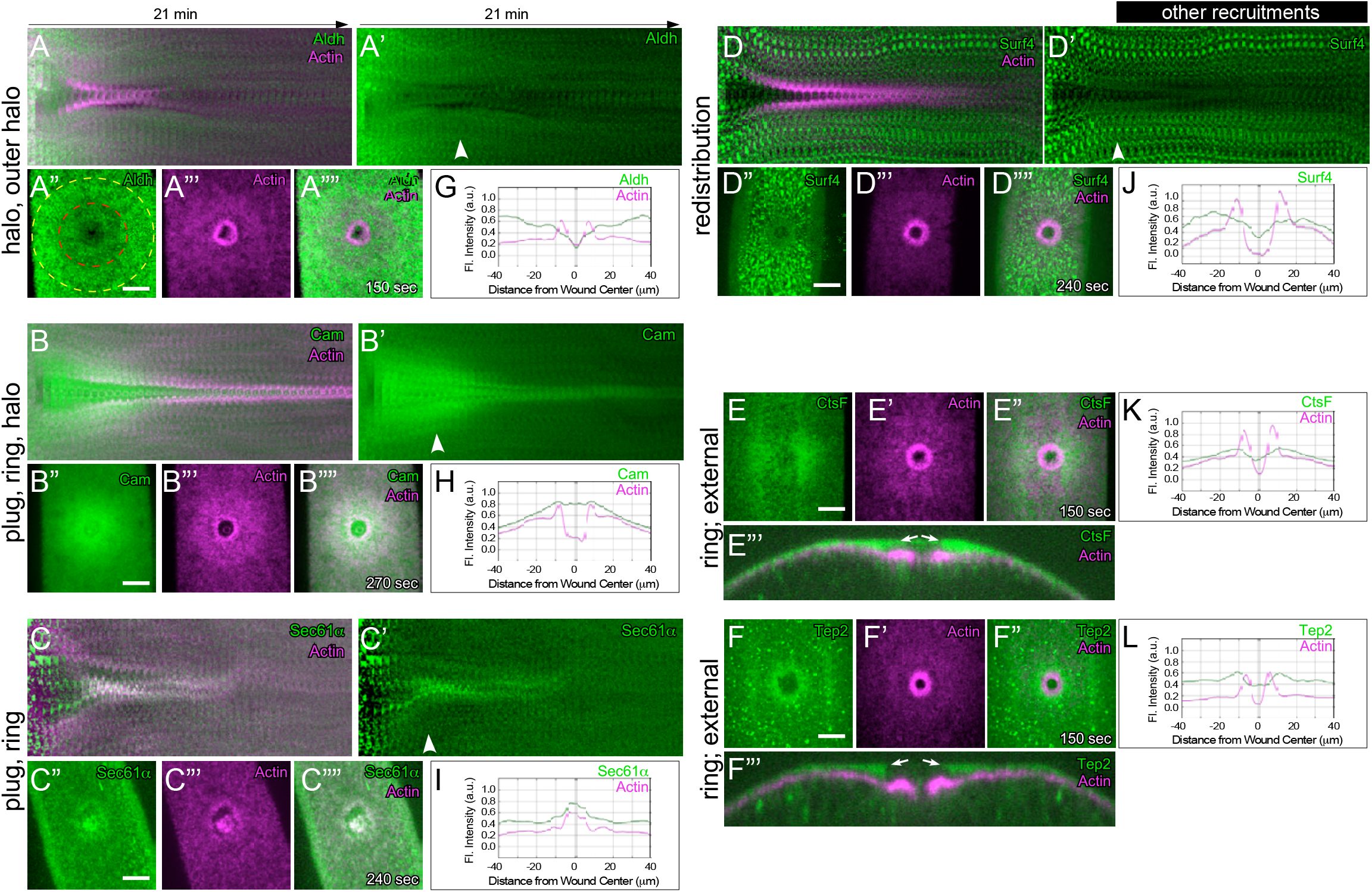
Proteins exhibiting other recruitment patterns to cell wounds. (A-D””) Confocal max projection images of embryos expressing a fluorescent actin reporter (sGMCA or sStMCA) and the following fluorescently-tagged proteins: Aldh (A-A””), Cam (B-B””), Sec61α (C-C””), or Surf4 (D-D””), at the time points indicated. (A-D’) Kymographs across the wound area in A-D””, respectively. White arrows indicate the time point of the XY views shown in A”-D””. (A”-D””) XY views of the time point indicated across the wound area in A-D””, respectively. The red dashed circle in A” denotes actin halo region and the yellow dashed circle denotes the outer halo region. (E-F’”) Confocal max projection images of embryos expressing a fluorescent actin reporter (sGMCA or sStMCA) and the following fluorescently-tagged proteins: Ctsf (E-E’”) or Tep2 (F-F’”), at the time points indicated. (E-F”) XY views of the time point indicated across the wound area in E-F”, respectively. (E’”-F’”) Orthogonal view of E-F”, showing external recruitment of proteins around the wound site. (G-L) Fluorescence intensity (arbitrary units) profiles across the wound area over time for the images shown in (A””-D””, E-F”). Scale bars: 20μm.

### Rab family GTPases are recruited to cell wounds with different temporal and spatial patterns

In addition to the proteins just described, we identified the recruitment of a large number of Rab family GTPases to cell wounds. Rab GTPases are a large conserved family of small GTPases that regulate nearly all aspects of intracellular membrane trafficking from vesicle formation to vesicle movement along cytoskeleton networks and membrane fusion (59–67). Disrupted Rab GTPase function is associated with and/or causative of a wide range of diseases, including neurodegenerative and immune disorders (64, 66, 68–70). Rab GTPases are expressed in many different dynamic patterns and levels during development (71–74). Individual Rab GTPases localize to different membrane compartments through their stable C-terminal prenylation, where they recruit various downstream effectors to affect membrane lipid composition, identity, integrity, and function (64–66, 73). Not surprisingly, since membrane and cytoskeleton are highly coordinated during many cellular processes, depletion of Rab GTPase function has also been shown to disrupt the integrity of the cytoskeleton, including inhibiting actin assembly and modulating filament assembly at specific membrane locations (62, 63, 67, 75). Important for cell wound repair, these functions can affect wound closure and cell cortex remodeling (4, 76, 77).

Of the 27 Rab family GTPases expressed in the *Drosophila* syncytial embryo, we identified 15 Rabs that are recruited to cell wounds (Fig 15; Fig 16; Table 1). These Rab GTPases exhibit different spatial recruitment patterns and there are representatives for five of the seven broad categories of recruitment patterns: plug – Rab1, Rab2, Rab18 (Fig 15B-C’, 15I-J; Fig 16D-D’, 16L); actin ring – Rab6, Rab7, Rab8, Rab9, Rab10, Rab21, Rab40 (Fig 15F-H’, 15M-O; Fig 16A-B’, 16E-E’, 16H-H’, 16I-J, 16M, 16P); actin ring + actin halo – Rab5, Rab11, Rab35 (Fig 15E-E’, 15L; Fig 16C-C’, 16G-G’, 16K, 16O); multiple (inner ring + actin ring + actin halo): Rab23 (Fig 16F-F’, 16N); and Other: Rab4 (Fig 15D-D’, 15K). While the recruitment many of these Rabs peak early in the repair process, some of them are much later (cf. Rab4, Rab5, Rab9), indicating roles in the remodeling phase of the repair process (Fig 15D-E’, 15L-M; Fig 16A-A’, 16I). The wide spectrum of similarities and differences in their spatial and temporal recruitment suggests that each Rab is likely required for both specific roles and for overlapping ones during the cell wound repair process.

**Figure 15.**
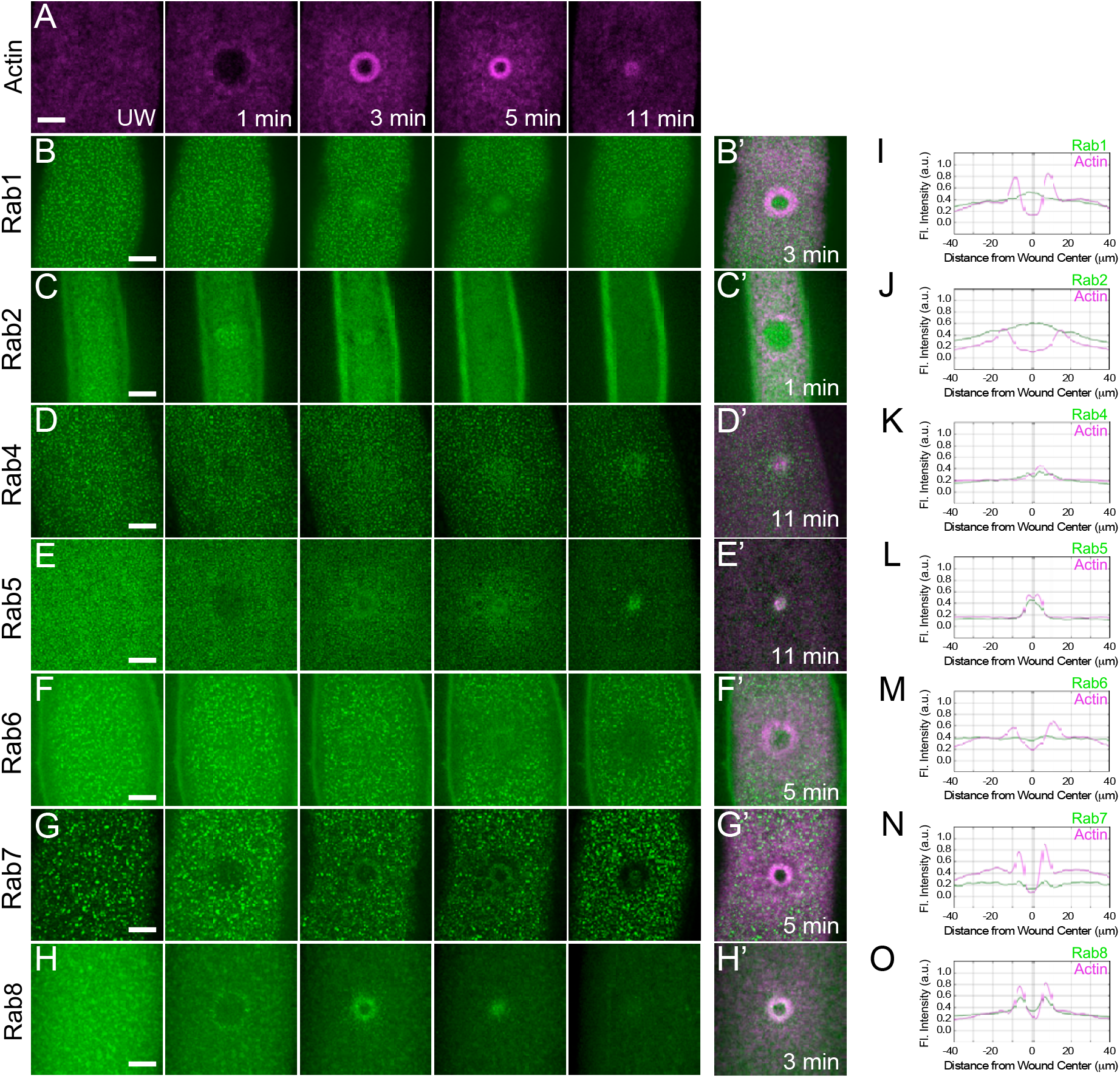
Recruitment patterns of Rab family GTPases to cell wounds. (A-H’) Confocal max XY projection images of embryos expressing a fluorescent actin reporter (sGMCA or sStMCA) (A) and the following fluorescently-tagged Rab GTPase proteins: Rab1 (B-B’), Rab2 (C-C’), Rab4 (D-D’), Rab5 (E-E’), Rab6 (F-F’), Rab7 (G-G’), or Rab8 (H-H’), at the time points indicated. (I-O) Fluorescence intensity (arbitrary units) profiles across the wound area over time for the images shown in (B’-H’). Scale bars: 20μm.

**Figure 16.**
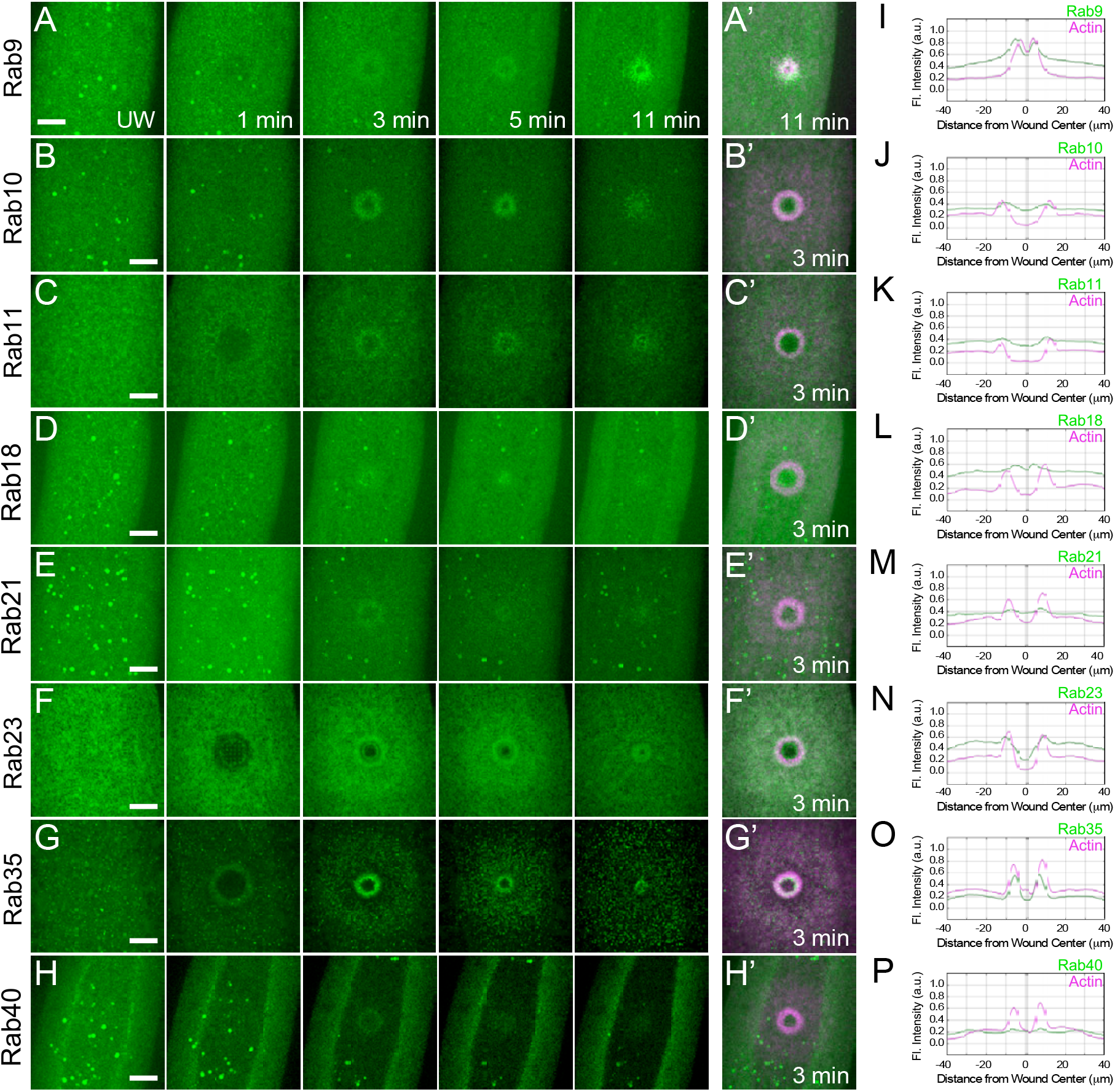
Recruitment patterns of Rab family GTPases to cell wounds (2). (A-P’) Confocal max XY projection images of embryos expressing a fluorescent actin reporter (sGMCA or sStMCA) and the following fluorescently-tagged Rab GTPase proteins: Rab9 (A-A’), Rab10 (B-B’), Rab11 (C-C’), Rab18 (D-D’), Rab21 (E-E’), Rab23 (F-F’), Rab35 (G-G’) or Rab40 (H-H’), at the time points indicated. (I-P) Fluorescence intensity (arbitrary units) profiles across the wound area over time for the images shown in (A’-H’). Scale bars: 20μm.

### Knockdown of individual Rab family GTPases results in distinct impaired cell wound repair phenotypes

We next examined the effects of removing 12 of the 15 recruited Rab family GTPases on cell wound repair. We generated knockdown embryos by one of 3 methods depending on the availability and efficacy of available reagents: 1) GFP-Rab fusion proteins were knocked down in the female germline by expressing a GFP RNAi construct with the GAL4-UAS system (67, 78, 79); 2) Traditional RNAi knockdowns were generated by expressing Rab GTPases RNAi constructs in the female germline using the GAL4-UAS system (80); or 3) in the case of Rab23, a homozygous null mutant exists (81) (Fig 17A) (see Methods). We were unable to examine Rab2, Rab4, or Rab5: Rab2 knockdowns do not produce embryos for analysis and the existing reagents for Rab4 and Rab5 did not generate sufficient knockdown for analysis. We then observed actin dynamics following laser wounding using a fluorescent actin reporter (sStMCA) (Fig 17, Fig 18). In all cases, the wounded knockdown embryos exhibited actin cytoskeleton disruptions at various post-initiation steps of the cell wound repair process, including wound over-expansion (Fig 17P), delayed/altered rates of wound contraction (Fig 17Q), and/or aberrant actin dynamics (Fig 18M-N). Examples of these phenotypes are described below.

**Figure 17.**
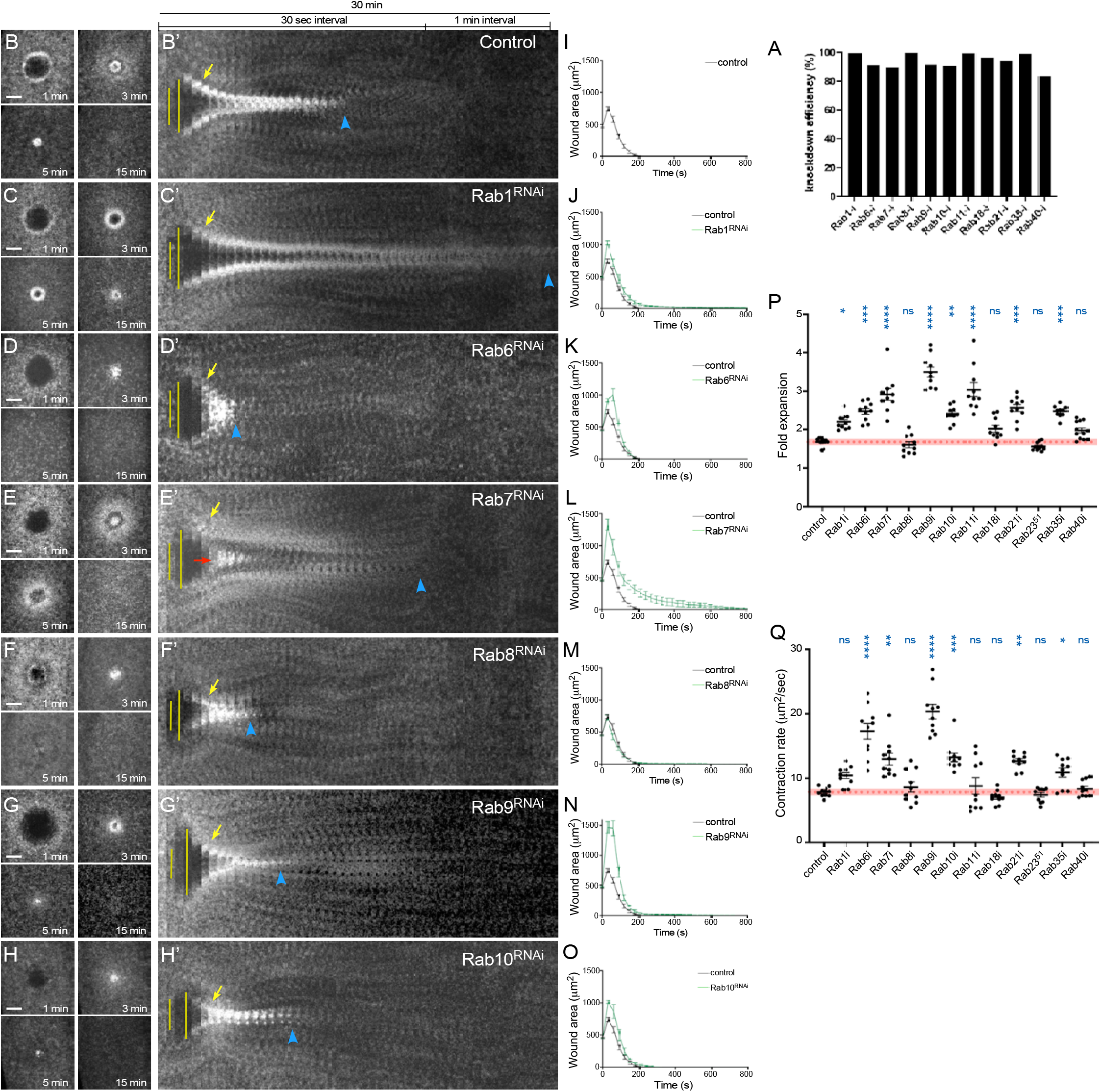
Knockdown of individual Rab family GTPases result in distinct cell wound repair phenotypes. (A) Quantification of knockdown efficiencies for Rab RNAi embryos under our imaging conditions determined by qPCR or western blot analysis. (Rab23 ^51^ is an amorphic mutant allele (79)). (B-H’) Confocal max projection images of wounds generated in embryos expressing an actin marker (sStMCA) in control (GFP RNAi; B), Rab1 RNAi (C), Rab6 RNAi (D), Rab7 RNAi (E), Rab8 RNAi (F), Rab9 RNAi (G), and Rab10 RNAi (H). (B’-H’) Kymographs across the wound area depicted in B-H, respectively. Wound expansion is highlighted by yellow lines. Actin recruitment to the actomyosin ring is indicated by yellow arrows. Actomyosin ring disassembly (or lack thereof) is indicated by blue arrowheads. Actin accumulation internal to the wound is indicated by red arrows. Scale bars: 20μm. (I-O) Quantification of the wound area over time for knockdowns shown in (B-H), respectively. (O-Q) Quantification of wound dynamics in Rab family GTPase knockdowns. Quantification of fold wound expansion (O) and wound contraction rate (P). Black line and error bars represent mean ± SEM. Red dotted line and square represent mean ± 95% CI from control. Kruskal-Wallis test was performed: * *p*<0.05, ** *p*<0.01, *** *p*<0.001, **** *p*<0.0001, ns is not significant.

**Figure 18.**
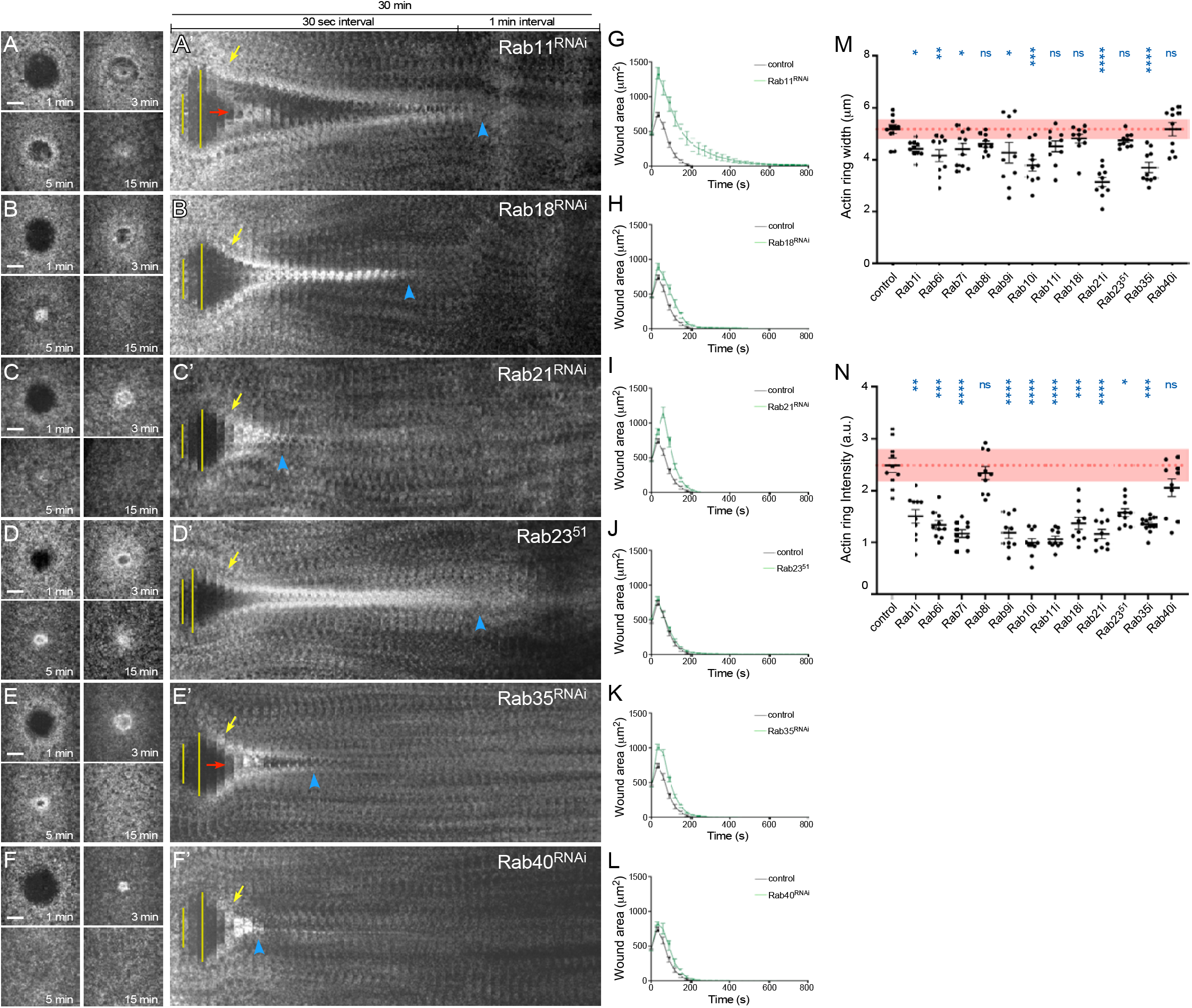
Knockdown of individual Rab family GTPases result in distinct cell wound repair phenotypes (2). (A-F’) Confocal max projection images of wounds generated in embryos expressing an actin marker (s StMCA) in Rab11 RNAi (A), Rab18 RNAi (B), Rab21 RNAi (C), *Rab23^51^* mutant (D), Rab35 RNAi (E), and Rab40 RNAi (F). (A’-F’) Kymographs across the wound area depicted in A -F, respectively. Wound expansion is highlighted by yellow lines. Actin recruitment to the actomyosin ring is indicated by yellow arrows. Actomyosin ring disassembly (or lack thereof) is indicated by blue arrowheads. Actin accumulation internal to the wound is indicated by red arrows. Scale bars: 20μm. (G-L) Quantification of the wound area over time for knockdowns shown in (A -F), respectively. Black line and error bars represent mean ± SEM. (M-N) Quantification of actin ring dynamics in Rab family GTPase knockdowns. Quantification of actin ring width (M) and actin ring intensity (N). Black line and error bars represent mean ± SEM. Red dotted line and square represent mean ± 95% CI from control. Kruskal-Wallis test was performed: * *p*<0.05, ** *p*<0.01, *** *p*<0.001, **** *p*<0.0001, ns is not significant.

In control embryos, a robust actomyosin ring forms within 1 minute post-wounding, progressively contracts to close the wound by approximately 13 minutes after injury, and is then disassembled (Fig 17B-B’, 17I, 17P-Q; Fig 18M-N). While knockdowns of all of the Rabs assemble an actomyosin ring, these are not as robust as in controls, and those generated in Rab6, Rab8, Rab9, Rab10, Rab21, Rab35, and Rab40 knockdown backgrounds are prematurely disassembled (Fig 17D-D’, 17F-H’, 17K, 17M-Q; Fig 18C-C’, 18E-F’, 18I, 18K-N). In the case of Rab35 knockdowns, the actomyosin ring is disassembled prior to wound closure (Fig 18E-E’, 18K). In contrast, the actomyosin rings generated in Rab1 and Rab23 persist much longer than in controls, suggesting a role in the remodeling phase of the repair process (Fig 17C-C’; Fig 18D-D’). Interestingly, knockdowns of Rab7 and Rab11 exhibit similar repair phenotypes with very delayed wound closure and an accumulation of action in the interior region of the wounds (Fig 17E-E’; Fig18A-A’). The syncytial embryo cortex is under tension such that when an embryo is wounded, the release of this resting tension leads to the wound area expansion through a recoiling of the cell cortex. Several of the Rabs exhibit wound over-expansion (Fig 17P), suggesting that they play a role in cell cortex partitioning with respect to tension. The breadth of cell wound repair phenotypes observed among the Rabs are consistent with their highly dynamic spatiotemporal recruitments to cell wounds, and their likelihood of playing roles alone and together for many aspects of the repair process.

## Discussion

Single cell wound repair is required extensively throughout normal everyday life, as well as in response to damage caused by diseases and cancers (1, 2, 4, 5, 23). Single cell wound repair is a complex process that requires tight spatial and temporal coordination of numerous molecules and machineries such as those controlling actin polymerization, actin/MT crosstalk, and trafficking of membrane. Here we identify 129 proteins that are recruited to cell wounds, providing a more comprehensive view of the cellular response to injuries, uncovering new entry points for examining specific steps in the cell wound repair process, and uncovering interesting aspects of biology (Fig 19). For example, we identified both the light and heavy chains of clathrin, but interestingly, their spatial recruitment patterns place them in separate categories: the heavy chain (CHC) is recruited to the halo region (Fig 12B-B””, H), whereas the light chain (Clc) is recruited initially to the halo region, but then concentrates in a dense ring overlapping the actin ring (Fig 13Ci-viii, I-I’). These differences in spatial localization could limit the regions and timing where a clathrin complex would be functional, and/or may indicate unexplored non-canonical roles for the individual components. In addition, Osbp, a lipid-transfer protein (82, 83), and Rtnl1, an ER-shaping reticulon-family protein (84, 85), are recruited to a ring immediately interior to the actin ring (Fig 3H-H””, Q; Fig 13Diii-v, J). This domain coincides with the interface between the transient membrane plug and the plasma membrane, raising the possibility that Osbp and Rtnl1 delineate sites of ER–PM fusion.

**Figure 19.**
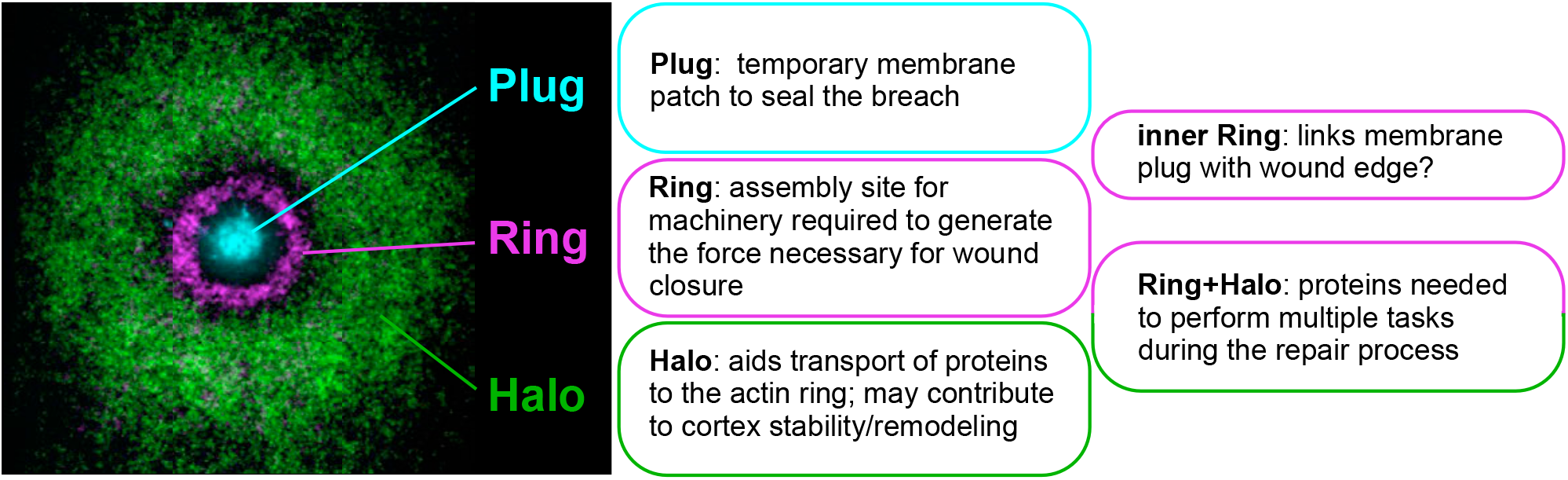
Summary of protein recruitments to and potential roles at cell wounds. (left) Image depicts the three major regions of protein recruitment to cell wounds (Plug, Ring, Halo) and (right) a description of their roles in the repair process.

The relatively recent introduction of the Drosophila model system in the study of cell wound repair has allowed global screening approaches to be used to identify molecules involved in cell wound repair (26). We previously performed a microarray screen to identify molecules involved in the transcriptional initiation of cell wound repair and surprisingly found that it is translation that is initially required for robust actomyosin ring formation and efficient cell wound repair, whereas transcription is required for subsequent steps in the Drosophila model (26). These findings suggest that pre-existing mRNAs and proteins can mediate the immediate cellular response to wounding and function during the early stages of repair. Consistent with this model, our genetic screen of fluorescently tagged proteins identified 129 molecules that are recruited to cell wounds, indicating that many proteins undergo rapid spatial and temporal redistribution upon wounding. This recruitment can be carried out by the rapid relocation of pre-existing protein pools or through the rapid translation of existing mRNAs. Notably, only six genes we identified here—Aldh, dmGlut, nebu, Pi3k21B, ReepB, Sec61α, and sstn—overlap with our previous microarray positives where they exhibited decreased transcript levels upon wounding. Together, our results support a model in which immediate wound repair is driven primarily by the redistribution of pre-existing proteins and the utilization of existing mRNA pools, whereas transcriptional responses become important during later stages of cell wound repair.

A recent proteome-scale screen in budding yeast identified 80 cell wound repair protein candidates through systematic analysis of fluorescently-tagged proteins (17). Among these 80 candidates, 43 have homologs in Drosophila, and 22 were examined in our genetic screen. Only 4 of these 22 proteins overlapped with our positive hits (RhoGEF2, WASp, Epsin, twinstar). In this case, the wounds induced in the budding yeast screen are small (0.5 µm diameter), with endocytosis and exocytosis thought to mediate the rapid removal and restoration of such small plasma membrane lesions (1–5). Larger wounds, such as those generated in our Drosophila model (20 µm diameter) require actomyosin ring assembly and contraction to physically close the damaged cell cortex (8, 10). Thus, the limited overlap between the budding yeast and Drosophila screens likely reflects multiple, context-dependent repair mechanism differences rather than a lack of conservation. Together, these genetic and proteomic approaches support the idea that cells choose distinct repair machineries according to various types of damage and are providing new insights into the mechanisms by which cells sense, respond to, and repair wounds of varying sizes and types.

Among the 129 proteins identified here, one family of proteins is highlighted: Rab GTPases, master regulators of membrane trafficking. Consistent with a trafficking role for Rabs in the repair process, we identified two cathepsins—CtsF and CtsL1—that are known to rely on Rab family GTPases for their trafficking and endolysosomal maturation (86–88). Cell wound repair can also use endo- and exocytosis pathways to remove the damaged lesion from the cell (1–5). Indeed, previous studies showed that Rab3, Rab10, and Rab11 are required for lysosome positioning near the wound to reseal the cell damage by pore-forming toxins (89–91), and Rab5 and Rab11 are required for cell survival from pore-forming toxins (92). However, our Drosophila model uses actomyosin ring contraction for wound closure, not endo-/exo-cytosis. We find that 15 out of 33 Drosophila Rab GTPases are recruited to the wound and exhibit distinct recruitment patterns and knockdown phenotypes. Interestingly, all of these Rab GTPase knockdowns affect actin dynamics during the repair process. Previous studies have uncovered emerging functions of Rab GTPases in the control of actin cytoskeletal organization and dynamics, suggesting roles beyond their canonical functions in vesicle trafficking (93–95). For example, Rab40b-Cullin5 complex ubiquitinates the actin bundling protein, EPLIN, during cell migration (94). Rab11 organizes Myosin II to regulate actomyosin contractility for apical constriction during tissue invagination (93). We recently found that Rab35 regulates actin oxidation and reduction through Mical and SelR to organize actomyosin ring formation and function during cell wound repair (95). In addition, several

Rab GTPases are canonically part of the same pathway, yet their recruitment patterns and knockdown phenotypes in the context of cell wound repair are not identical. While Rab1, Rab2, Rab6, and Rab10 function together during the secretory pathway (96–99), Rab1 and Rab2 are recruited to the plug region, but Rab6 and Rab10 are recruited to the ring region. Rab1 exhibits prolonged actin accumulation, whereas Rab6 and Rab10 exhibit premature actin ring disassembly during cell wound repair. Thus, cell wound repair requires non-canonical functions of these Rab GTPases to regulate different aspects of actin dynamics during the repair processes. Future investigations into how individual Rab GTPases regulate distinct aspects of actin dynamics during cell wound repair will provide new insights into the noncanonical functions of Rab GTPases in actin cytoskeleton regulation across diverse biological contexts.

In summary, we successfully identified 129 proteins that are recruited to cell wounds, many of which have not previously been examined in the context of cell wound repair. The functional classification of these proteins provides a valuable framework for dissecting the molecular machineries and mechanisms that govern distinct steps of the repair process, including damage sensing, membrane plug formation, actomyosin ring assembly and contraction, and cell cortex remodeling after wound closure. In addition, our findings establish a foundation for future mechanistic studies. Notably, our investigation of Rab GTPases highlighted their emerging roles in regulating actin dynamics within cells. Given that defective wound repair contributes to the pathogenesis of numerous diseases, including muscular dystrophies, neurodegenerative disorders, diabetes-associated tissue injury, and cancer progression, the proteins identified in this study may contribute to understanding cellular resilience and tissue homeostasis. Together, this work advances our understanding of the complex and highly coordinated processes that restore cell integrity after wounding and provides a foundation for uncovering new principles of cell wound repair, with important ramifications for therapeutic targets, potential treatments, and/or refinements to existing treatment modalities.

## Acknowledgements

We thank Tony Cooke for his microscopy wizardry, and former Parkhurst lab members for technical help in different parts of the project. We thank Lynn Cooley, FlyBase, the Bloomington Stock Center, the Kyoto Stock Center, the Harvard Transgenic RNAi Project, the Vienna Drosophila Stock Center, and the Fred Hutch/Leica Center of Excellence for advice, microscopes, flies, and other reagents used in this study. This work was supported by NIH grants R35GM161275 and R01GM111635 and the Mark Groudine Chair for Outstanding Achievements in Science and Service (to SMP).

## Author Contributions

All authors contributed to the design of the experiments, performed experiments, analyzed data, and performed the morphometric analyses. MN and SMP wrote the manuscript with input from all authors.

## Materials and Methods

Reagents used in this study are described in Table S1.

### Fly stocks and genetics

Flies were cultured and crossed at 25°C on yeast-cornmeal-molasses-malt medium. Flies used in this study are described in Table S1 and Table S2. All fly stocks were treated with tetracycline, then tested by PCR to ensure that they did not have Wolbachia.

To knockdown Rab genes, we used one of 3 methods depending on the availability and efficacy of available reagents: 1) GFP-Rab fusion proteins were knocked down in the female germline by expressing a GFP RNAi construct with the GAL4-UAS system for Rab1, Rab6, Rab7, Rab9, Rab10, Rab11, Rab18, Rab21, and Rab40 (78, 79, 100); 2) Traditional RNAi knockdowns were generated by expressing Rab GTPases RNAi constructs in the female germline using the GAL4-UAS system for Rab4, Rab5, Rab8, and Rab35 (80); or 3) a previously described Rab23 homozygous mutant was used (81). RNAi and fluorescent-tagged protein lines were driven maternally using the GAL4-UAS system with P{matalpha4-GAL-VP16}V37 (Bloomington #7063 or #7062) or P{w[+mC]=GAL4::VP16-nos.UTR}MVD1 (Bloomington #4937). Knockdown efficiency was tested by qPCR or western blot analyses.

An actin reporter, sGMCA (spaghetti squash driven, moesin-alpha-helical-coiled and actin binding site fused to GFP) reporter (101) or the mScarlet-i (sStMCA (27)) or mCherry (sChMCA (8)) fluorescent equivalents were used to follow wound repair dynamics of the cortical cytoskeleton. Mutant analyses were performed at least twice from independent genetic crosses and ≥10 embryos were examined unless otherwise noted. Images representing the average phenotype were selected for figures.

### Generation of fluorescently tagged transgenic flies

To generate sqh-sfGFP-CHC, the CHC ORF was amplified from BDGP clone LD43101 and fused 5’ to sfGFP. The resulting sfGFP-CHC fusion was cloned into pSqh5′+3′UTR (8) as a 5′ StuI-3′ XbaI fragment.

To generate sqh-StFP-Clc, the CHC ORF was amplified from BDGP clone GM02293 and fused 5’ to mScarlet-i (Addgene #85044). The resulting StFP-CHC fusion was cloned into pSqh5′+3′UTR as a 5′ StuI-3′ XbaI fragment.

To generate UASp-ChFP-Jbug, the Jbug ORF was amplified from BDGP clone RE40504 and fused 5’ to mCherry, The resulting ChFP-Jbug was cloned into pUASp as 5’ NotI-3’ XbaI fragment.

To generate sqh-StFP (mScarlet-i) -SelR, the SelR ORF was amplified from BDGP clone RE73235 and fused 5′ to StFP. The resulting StFP-SelR fusion was cloned into pSqh5′+3′UTR (7) as a 5′ StuI-3′ XbaI fragment.

To generate sqh-GFP-p120ctn, the p120ctn ORF was amplified from BDGP clone LD33274 and fused 5’ to GFP. The resulting GFP-CHC fusion was cloned into pSqh5′+3′UTR as a 5′ StuI-3′ XbaI fragment.

Constructs (500μg/ml) were injected along with the pTURBO helper plasmid (100μg/ml) into isogenic w1118 flies as previously described (102). Transgenics were scored by eye color and the insertions were mapped using standard genetic methods.

### Embryo handling and preparation

Nuclear cycle (NC) 4-6 *Drosophila* embryos were collected from 0-30 min at room temperature (22°C). Embryos were hand dechorionated, placed onto No. 1.5 coverslips coated with glue, and covered with Series 700 halocarbon oil (Halocarbon Products Corp).

### Laser wounding

All wounds were generated with a pulsed nitrogen N2 Micropoint laser (Andor Technology Ltd., Concord, MA, USA) tuned to 435 nm and focused on the cortical surface of the embryo. A region of interest was selected in the lateral midsection of the embryo and ablation was controlled by MetaMorph. On average, ablation time was less than 3s, and time-lapse imaging was initiated immediately. Occasionally, a faint grid pattern of fluorescent dots is visible at the center of wounds that arises from damage to the vitelline membrane that covers embryos.

### Microscopy

All imaging was performed at room temperature (22°C). A Zeiss Axioplan 200 microscope with an Apotome and digital camera was used for the initial screening. The following microscopes were used for the subsequent dynamic live imaging: 1) Revolution WD systems (Andor Technology Ltd., Concord, MA, USA) mounted on a Leica DMi8 (Leica Microsystems Inc., Buffalo Grove, IL, USA) with a 63x/1.4 NA objective lens and controlled by MetaMorph software. Images and videos were acquired with 488 nm and 561 nm, using an Andor iXon Ultra 897 or 888 EMCCD cameras (Andor Technology Ltd., Concord, MA, USA). 2) A Yokogawa CSU-X1 confocal spinning disk head mounted on a Nikon Eclipse Ti (Nikon Instruments, Melville NY,USA) with a 60x/1.4 NA objective lens and controlled by MetaMorph software. Images and videos were captured using 488nm and/or 561nm lasers using an Andor iXon Ultra 888 EMCCD camera (Andor Technology Ltd., Concord, MA, USA). All images for cell wound repair were 17-20 µm stacks/0.25 µm steps. For single color, images were acquired every 30 sec for 15 min and then every 60 sec for 25 min. For dual green and red colors, images were acquired every 30 sec for 30-40 min.

### Image processing, analysis, and quantification

All images were analyzed with Fiji (103). Measurements of wound area, wound expansion rate, wound contraction rate, actin ring intensity and width were performed as previously described (29, 30). Measurements of wound area were done manually. To generate xy kymographs, all time-lapse xy images were cropped to 5.8 µm x 94.9 µm and then each cropped image was lined up. For fluorescent line plots, the mean fluorescence profile intensities were calculated from 51 equally spaced radial profiles anchored at the center of the wound, swept from 0° to 180°. Radial profiles of diameter 301 pixels were used (24). The lines represent the averaged fluorescent intensity and gray area is the 95% confidence interval. Line profiles from the left to right correspond to the top to bottom of the images unless otherwise noted.

Quantification of the width and average intensity of actin ring, wound expansion, and closure rate was performed as follows: the width of actin ring was calculated with two measurements, the ferret diameters of the outer and inner edge of actin ring at 90 sec post-wounding. Using these measurements, the width of actin ring was calculated with (outer ferret diameter – inner ferret dimeter)/2. The average intensity of actin ring was calculated with two measurements. Instead of measuring ferret diameters, we measured area and integrated intensity in same regions as described in ring width. Using these measurements, the average intensity in the actin ring was calculated with (outer integrated intensity - inner integrated intensity)/(outer area - inner area). To calculate relative intensity for unwounded (UW) time point, average intensity at UW was measured with 50×50 pixels at the center of embryos and then averaged intensity of actin ring at each timepoint was divided by average intensity of UW. Wound expansion was calculated with max wound area/initial wound size. Closure rate was calculated with two time points, one is t_max_ that is the time of reaching maximum wound area, the other is t<half that is the time of reaching 50-35% size of max wound since the slope of wound area curve changes after t<half. Using these time points, average speed was calculated with (wound area at t_max_ – wound area at t<half)/t_max_-t<half.

### qPCR

Total RNA was obtained from 100 embryos (0-30 min old) using TRIzol (Invitrogen). 1 µg of total RNA was used for reverse transcription with the iScript™ gDNA Clear cDNA Synthesis Kit (Bio-Rad). RT-PCR analysis was performed using the iTaq™ Universal SYBR® Green Supermix (Bio-Rad) with two individual parent sets and two technical replicates on the CFX96TM Real Time PCR Detection System (Bio-Rad). RpL32 was used as a reference gene. The % knockdown was calculated using the ΔΔCq calculation method compared with control (vermilion knockdown). Same primer sets RpL32 was used in previously described (26).

### Western blots

To generate lysates, 0-30min embryos were homogenized in 2x sample buffer (1x sample buffer: 0.5M Tris pH6.8, 10% glycerol, 2% SDS, 5% 2-mercaptoethanol, 0.005% bromophenol blue), then centrifuged at 14K rpm for 1 minute at 4°C to remove cellular debris. Lysate samples were subjected to SDS-PAGE electrophoresis and then blotted to nitrocellulose membrane according to standard procedures. The following antibodies were used: anti-GFP mouse monoclonal (1:1000; Roche), with anti-actin (A2103; 1:2500; Sigma) as the loading control. Quantification of proteins on westerns was performed using ImageJ (National Institutes of Health, Bethesda, MD). A minimum of two biological replicates were performed for each western.

### Statistical analysis

All statistical analysis was done using Prism 8 (GraphPad, San Diego, CA). Gene knockdowns were compared to the appropriate control, and statistical significance was calculated using a Kruskal-Wallis test with p<0.05 considered significant.

